# Hawaiian Geothermal Fumaroles Contain Diverse and Novel Viruses

**DOI:** 10.64898/2026.04.06.716669

**Authors:** Pia Sen, Lausanne Lee Oliver, Kira S. Makarova, Yuri I. Wolf, Christina Pavloudi, Max Shlafstein, Jimmy H. Saw

**Author notes:** Co-Corresponding authors: Pia Sen & Jimmy H. Saw.

## Abstract

Microbial communities of geothermal habitats are central to understanding the evolution of life on Earth. Metagenomics has provided insight into the role of viruses in shaping microbial diversity of complex environments. However, identification of novel viruses is constrained by lack of marker genes and low nucleotide similarities between related viral taxa. While microbial and viral diversity have been explored in terrestrial hot springs and hydrothermal vent systems, other volcanic features remain underexplored. Fumaroles (steam vents) are geothermal features that heat groundwater with magma, releasing steam and volcanic gases such as CO2 and H2S. Comparatively physicochemically dynamic to hot springs, fumarole temperatures and gas emissions rapidly fluctuate with volcanic activity. Here, we describe viruses identified metagenomically from microbial mats hosted near basaltic fumaroles on the Big Island of Hawaìi. To our knowledge, this is the first systematic survey of fumarole viruses. Our utilization of a sensitive profile-based approach for identification reveals high viral diversity in fumaroles, resulting in estimation of two undescribed order-level clades of Caudoviricetes (tailed phages). Viral metabolic genes provide evidence of viral-mediated adaptation of microbes to fumarole conditions. We describe patterns of viral diversity that diverge from the Bank model of viral ecology, hinting at viral dispersal between biofilms and high viral richness and evenness. Lastly, we provide a description of the first terrestrial geothermal environment dominated by Microviridae, previously only described in viral communities of deep ocean hydrothermal vents. This study offers important findings for exploration of viral ecology in extreme environments.

## Introduction

Fumaroles (terrestrial steam vents) are the most common, yet least explored geothermal feature on Earth. Fumaroles are formed when steam and volcanic gases exit cracks in basalt deposits (Wall et al., 2015). Temperature and volcanic gas emissions fluctuate with volcanic activity and geological context (Prescott et al., 2022). Located on the volcanically active Big Island of Hawaìi, the East Rift Zone (ERZ) is a geologically active region in the vicinity of Hawaii Volcanoes National Park. Along Hawaii Route 130 and between the towns of Pahoa and Kalapana, there are many physicochemically diverse steam vents, which produce steam that is mainly composed of water. These fumaroles occur on two lava flows with a wide age range (64-400 years old) and host phototrophic biofilms dominated by *Cyanobacteria*, *Chloroflexota*, *Deinococcota*, and *Alphaproteobacteria* (Prescott et al., 2022). Previous investigation of these fumaroles reveals microbial succession patterns that are dependent on early colonizing phototrophs with Actinobacteria and Proteobacteria becoming more abundant in the biofilms as geothermal activity decreases in the system (King, 2003; Prescott et al., 2022). While microbial diversity of other geothermal systems is shaped by both interspecific competition and habitat filtering, microbial diversity of the steam vent biofilms is primarily explained by interspecific competition which results in an over dispersed microbial community. Functional annotation of microbial taxa from fumaroles suggests that photosynthesis, CO_2_ fixation, and central carbon metabolism are the primary biogeochemical processes in the biofilms. The presence of taxa associated with nitrogen and sulfur cycling indicated these processes may be relevant to biofilm biogeochemical cycling as well. Metals and minerals are made bioavailable in fumaroles by microbial alteration of basaltic substrate, while CO_2_ and H_2_S emissions from the vent stimulate carbon fixation and sulfur oxidation (Wall et al., 2015).

Microbial viruses modulate functional microbial diversity through one of three main actions: top-down predation of microbial hosts, transduction, or metabolic rewiring during infection which contributes to metabolic reprogramming of hosts on an ecosystem-level scale (Tian et al., 2024; Zimmerman et al., 2020). The Bank model of viral ecology describes ecosystems where a fraction of viruses are active and abundant, while a myriad of other viruses are inactive and present in low abundances until a shift in environmental conditions (e.g. seasonal or diel intervals) giving rise to new dominant host-virus pairs as observed in the succession patterns of environmental viral populations (Breitbart & Rohwer, 2005; Ji et al., 2024; Liang et al., 2019). The Bank model predicts that a viral community high in vOTU richness will be low in vOTU evenness. The density dependent ‘Kill-the-Winner’ (KtW) paradigm describes virulent viruses infecting the most abundant or fastest growing hosts, while the ‘Piggyback-the-Winner’ paradigm describes temperate viruses increasing in abundance with rates of lysogeny (Hu et al., 2025). While relative abundance cannot be taken as confirmation of viral activity, it is a metric that provides a good indicator of active viral populations (Liang et al., 2019; Zhou et al., 2025).

Viral biogeography cannot be separated from the biogeography of their microbial hosts, but is controlled by an interplay of viral characteristics (life cycle, burst and virion size) and abiotic factors. Geothermal systems are good models for exploring viral biogeography because their microbial communities are distinct and endemic islands across a broad range of thermophily (Castelán-Sánchez et al., 2020; Langwig et al., 2025; Power et al., 2018; Salgado et al., 2025; Teske & Reysenbach, 2015; Zhou et al., 2022). Viral dispersal in geothermal systems is limited and subject to distance-decay and strong habitat clustering patterns, but only within hot spring fields (Castelán-Sánchez et al., 2020; Cheng et al., 2022; Hwang et al., 2023; Salgado et al., 2025; Thomas et al., 2021). Microbial mats (biofilms) in geothermal systems are low mobility environments with highly specific host-virus pairs (Castelán-Sánchez et al., 2020; Jarett et al., 2020; Salgado et al., 2025).

First, we seek to describe patterns of viral dispersal and diversity across fumarole microbial populations. While microbial dispersal is known to vary across physicochemical profiles, geothermal systems of hot springs and hydrothermal vents have consistently been described as viral and microbial “islands”, with limited movement between both geothermal sites and smaller ranges of specific geothermal features within the broader system (Langwig et al., 2025; Prescott et al., 2022; Salgado et al., 2025). In such systems with high viral endemism and low dispersal, highly specific host-virus pairing is observed (Salgado et al., 2025). We gain insight into dispersal dynamics with viral read mapping and calculations of vOTU relative abundance. We predict host-virus pairing by matching microbial CRISPR spacers to viral genomes. If we observe CRISPR-based evidence of abundant vOTUs in an ‘arms race’ with a specific microbial population (many spacer matches to a single vOTU) we expect highly specific host-virus pairing and that there will be corresponding fluctuations in abundances of these viral and microbial populations across samples (Castledine et al., 2022). This would reflect lytic fumarole viral populations following the Bank model and adhering to the ‘Kill-the-Winner’ hypothesis, which describes a boom-bust dynamic where viruses target the dominant microbial population, triggering a proportional increase in vOTU abundance as viruses and their hosts engage in a coevolutionary arms race (Xue & Goldenfeld, 2017). This would reflect lytic fumarole viral populations following the Bank model and adhering to the ‘Kill-the-Winner’ hypothesis, which describes a boom-bust dynamic where viruses target the dominant microbial population, triggering a proportional increase in vOTU abundance as viruses and their hosts engage in a coevolutionary arms race (Bucher & Czyż, 2024). Third, we hypothesize that the viruses within the fumarole biofilms will contain genomic evidence of viral capability to facilitate microbial adaptation in a dynamic and extreme environment. We predict that viruses will contain auxiliary metabolic genes that could increase host fitness in volcanic systems.

In this study, we test these hypotheses with metagenomic identification and characterization of viruses in Pahoa fumarole biofilms from three different geothermal locations on the Big Island of Hawaìi (steam wall and steam pit within the East Rift Zone (ERZ), and the Big Ell cave in the Kilauea caldera). We use a sensitive and profile-based approach to identify and classify vOTUs, many of which are distantly related to known viruses. Further, we characterize viral genomes to identify shared predicted metabolic traits with viruses previously described in other early Earth analog geothermal environments. We find that ERZ fumaroles are a unique system for viral ecology, and demonstrate high viral dispersal and phylogenetic novelty. The presence of auxiliary metabolic gene (AMG) candidates demonstrates potential for viral influence on host metabolism. Our study provides insight into the diversification of early life and patterns of viral and microbial succession (Godin et al., 2023; Hadland et al., 2024).

## Results

### Sampling and retrieval of viruses from metagenome

Biofilm samples from ERZ fumaroles are described in previous studies (Prescott et al., 2022; J. H. Saw et al., 2026). Briefly, biofilms that were sampled were several centimeters thick and grew on the surface of volcanic rock, directly where steam was being released. These biofilms do not exhibit clear segmentation of layers as observed in some microbial mats, and display a variety of phenotypes and colors (J. H. Saw et al., 2026). Most Gloeobacter-dominated biofilms are purple with patches of dark green, while other biofilms are green, yellow, brown, and grey. While some biofilms have a visible extracellular polysaccharide matrix (EPS) layer and appear wet, other biofilms are dry or otherwise textured in appearance. While the steam pit and steam wall are separated by ∼100 meters, the Big Ell cave in the Kilauea Caldera is ∼35 km from the steam vents along Hawaii Route 130 (J. H. Saw et al., 2026).

WGS raw sequences from related biofilm features were co-assembled for related features (20 metagenomes from the 46 collected samples) with the ViralAssembly module of metaviralSPAdes, producing 400.15 Mbp (12,253 contigs) of assembled sequence data from the metaviralSPAdes pipeline (Antipov et al., 2020; J. H. Saw et al., 2026). 112.97 Mbp of the assembled sequence data was generated from Big Ell cave in the Kilauea Caldera (Hawaii Volcanoes National Park) with 17.87 Mbp (452 contigs) of the sequence data from geothermal biofilm features within the cave, and 95.11 Mbp (3803 contigs) of sequence data from adjacent non-geothermal volcanic soil samples. 243.97 Mbp (6596 contigs) of sequence data was assembled from exposed steam wall biofilms, and 43.21 Mbp (1402 contigs) of sequence data was assembled from the steam vents along Hawaii Route 130. From these assembled contigs, the ViralVerify module of metaviralSPAdes identified 754 contigs (6.33 Mbp) as either confidently (n=76) or uncertainly (n=678) viral (Supplemental Material S1). These contigs were then used in downstream analysis. VirSorter2 (Guo et al., 2021) identified 119.46 Mbp (n=16,002) of candidate viral contigs. From there, the VirSorter2 contigs (annotated with DRAM-V) were filtered for PFAM clan identifiers for the capsid genes (Phage-Coat: CL0373 and Inovirus-Coat: CL0371) producing 14.29 Mbp of candidate viruses (802 contigs) which were used for downstream analysis.

The accuracy of VirSorter2 filtering by PFAM clan was assessed by screening the remaining 15200 contigs identified as candidate viruses by VirSorter2 for the terminase large subunit gene (TerL) or another packaging gene (Supplemental Material S2). 320 contigs (∼2% of filtered VirSorter2 output) contained TerL and were manually assessed for viral identity, with upstream and downstream regions searched for other viral components. Of these contigs, 116 (0.76% of filtered contigs) were determined to be viral (containing both head and packaging genes). With geNomad (Camargo et al., 2024) we identified 3.41 Mbp (n=107) viral candidates (Supplemental Material S3). When final viral designations were made and vOTUs were generated, 24 vOTUs were identified by all of the tools used: VirSorter2, ViralVerify, and geNomad (Supplemental Figure 01). 15 vOTUs were identified by both VirSorter2 and ViralVerify, and a single vOTU was identified by both ViralVerify and geNomad. The majority of vOTUs (n=313) were identified from VirSorter2. 6 vOTUs were only identified by ViralVerify, and 5 vOTUs were identified only by geNomad.

### Remote homology-based identification and network analysis reveals diversifying vOTUs

569 viral contiguous sequences (contigs) were identified from 20 fumarole sample metagenomes (Supplemental Material S4). From these contigs, we performed clustering by community standards (Roux et al., 2019) and sorted viral sequences into 383 species-rank level vOTUs (viral populations). 51% of identified fumarole vOTUs are identified as Caudoviricetes (195 vOTUs), indicating that microbial viruses make up a significant portion of the metagenomically retrieved dsDNA viruses in fumarole biofilms. 46% of vOTUs (180 vOTUs) cannot be placed into realms by vConTACT3 v.3.0 despite verification of structural viral genes on each contig. These unclassified vOTUs may have ecological significance, making up at least 10% of viral relative abundance in the majority (17 out of 20) of samples (Figure 01).

**Figure 01.**
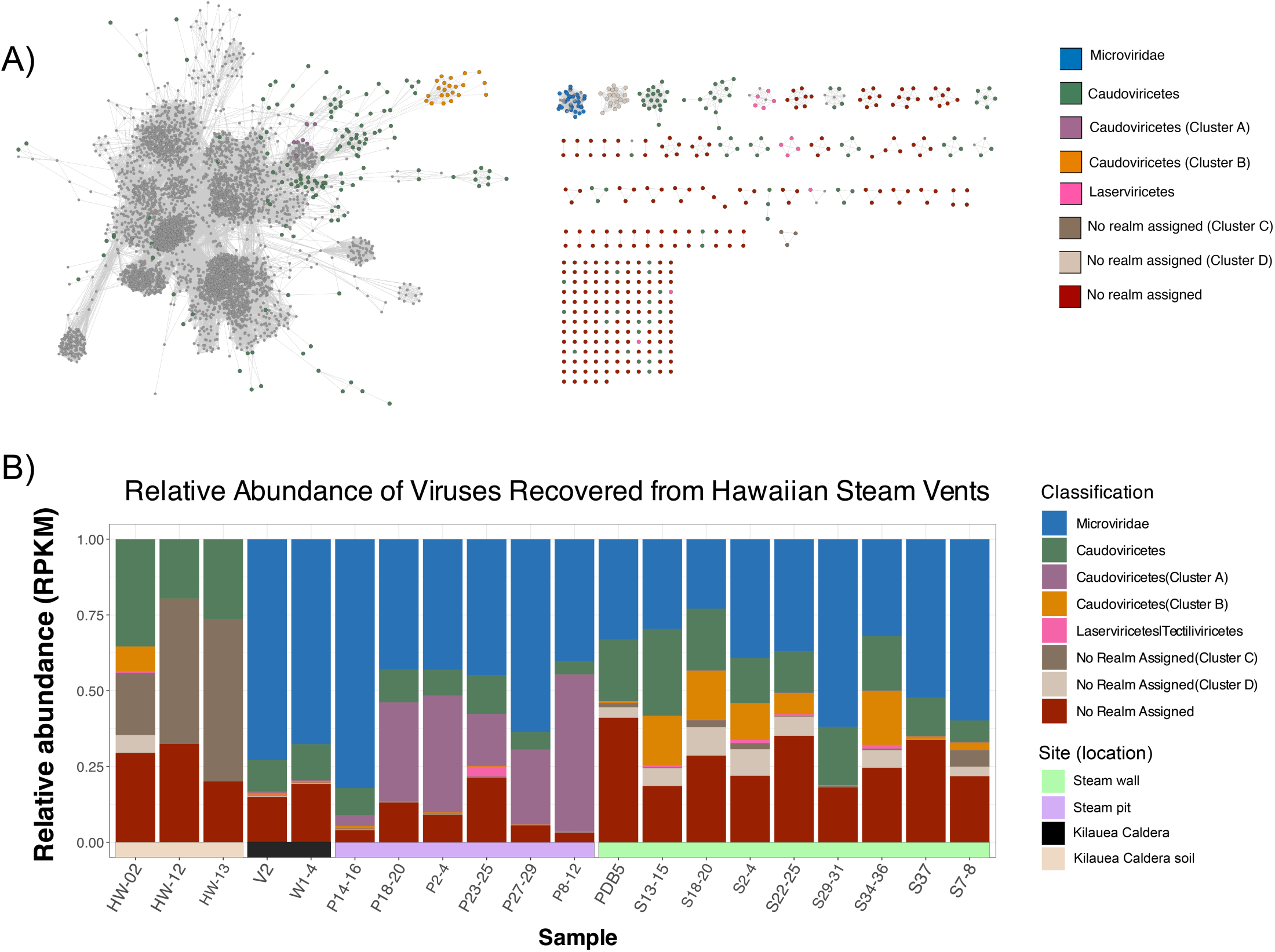
**A)** Gene-sharing network of confidently identified fumarole viral contigs visualized in Cytoscape. Nodes represent full viral sequences. Grey nodes are RefSeq viruses from the vConTACT3 v220 database, while colored nodes are fumarole viruses, grouped by vConTACT3 taxonomic predictions. Categories indicate the lowest confident taxonomic level assigned to each cluster; highly abundant clusters are colored independently of taxonomy. Edges represent the strength of gene-sharing between viruses, determined with protein clusters. All shown clusters have high clustering coefficients (>0.85), as calculated using the Cytoscape NetworkAnalyzer plugin. **B)** Relative abundance of identified viruses in all 20 samples, colored by cluster classification. Relative abundance (RPKM) was calculated for each individual viral contig and grouped by the taxonomic designation of clusters for visualization. Clusters found in notable abundance were colored and denoted independently, regardless of predicted taxonomy. Relative abundance of vOTUs used for statistical analysis is available as Supplemental Material S6.

Of the 180 vOTUs which could not be placed into a realm with network analysis, 100 of the ‘no realm’ vOTUs were missed by geNomad and filtered out in the VirSorter2 pipeline (Tian et al., 2024) during the CheckV step (Nayfach et al., 2021) suggesting stringent pipelines may miss some of the ecologically significant and highly divergent viruses (Camargo et al., 2024; Liang et al., 2019). The vOTUs were placed into genus-rank viral clusters (VCs) established by vConTACT3 v.3.0 and were visualized in Cytoscape (v3.10.2) with and without all viruses from NCBI RefSeq (Goldfarb et al., 2024). Clusters were filtered for quality by topological and clustering coefficients (>0.85). This network yielded 230 distinct topological groups, including subclusters, nested subclusters, and singletons within the network (Figure 01). Of these topological formations, 20 VCs form distinct subclusters (clusters of viral genomes with three or more genomes without links to the rest of the network). Multiple distinct viral populations (vOTUs) within a single VC provides evidence of genomic variation and diversification within the genus (Bolduc et al., 2025). High ‘Closeness Centrality’ values support the relationships observed within network clusters and prevent incomplete viral genomes from inflating network representation of genomic distance between vOTUs (Nayfach et al., 2021; Su et al., 2014). Resulting network clusters made up entirely of distinct fumarole vOTUs indicates that related viral populations share many genes with each other while still providing evidence of diversification between related vOTUs.

While 61% of Caudoviricetes clusters contain one vOTU, six clusters have between 8-24 different vOTUs in a single cluster (Figure 01). One highly abundant cluster, Caudoviricetes (Cluster B), makes up to 20% of viral relative abundance in three different biofilm samples from the steam wall. This cluster contains 20 unique vOTUs. The high abundance of related vOTUs is evidence of potential speciation occurring in abundant viral populations (Liang et al., 2019). There is also diversity observed within VCs of viruses that cannot be placed into a realm confidently by vConTACT3, providing tentative evidence of diversification within highly novel viral genera: out of the 21 VCs (three or more viral genomes within a cluster) which could not be realm placed, 11 of the ‘no realm’ filtered VCs contain more than a single vOTU (Supplemental Material S5). All of these 11 VCs are internally coherent with high connectivity measured by the NetworkAnalyzer plugin (Su et al., 2014).

### Phylogenetic estimation of viral novelty

#### Fumaroles contain new candidate Caudoviricetes orders

Out of the 196 vOTUs identified as Caudoviricetes by VContact3 assignment, 179 Caudoviricetes vOTUs contained a terminase large subunit (TerL) gene, and thus, were included in phylogenetic analysis with maximum likelihood (91% of Caudoviricetes vOTUs). Using 320 first neighbors of fumarole viruses from ICTV MSL40.v1, only 3% of these references could be placed into an existing order. Less than half (40%) of these references could be placed into previously identified Caudoviricetes families. The lack of classification capability for the reference viruses which are most closely related to fumarole vOTUs is evidence of fumarole viral novelty. Many fumarole viruses form distinct clades, with references often paraphyletic or sister taxa to fumarole viruses, suggesting that steam vent viruses expand the diversity of Caudoviricetes. The resulting maximum likelihood tree provides evidence of phylogenetic diversification and potentially speciation within Caudoviricetes clades. There is evidence of recent diversification within at least five clades that are almost entirely made up of fumarole viruses and contain 130 vOTUs (Figure 02). These species-rank vOTUs meet MIUViG standards necessary for taxonomic proposals (Roux et al., 2019).

**Figure 02.**
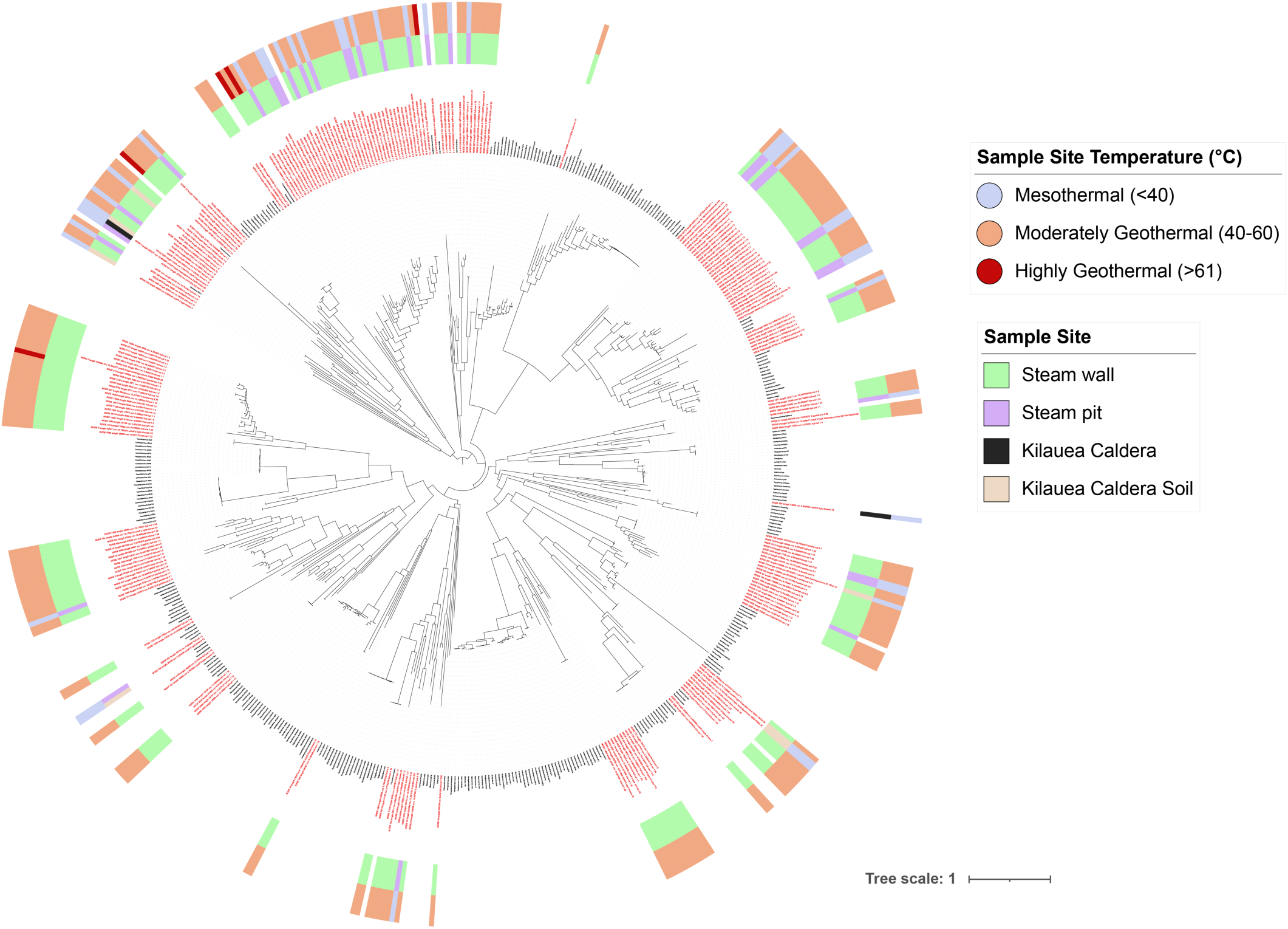
Maximum likelihood phylogenetic tree of Caudoviricetes terminase large subunit (TerL) amino acid gene sequences. Red node labels correspond to TerL sequences annotated in fumarole viral contigs. Black node labels correspond to RefSeq taxon which were determined to be first neighbors of fumarole viruses with Cytoscape plugin NetworkAnalyzer. Outmost ring refers to the temperature range of the sample from which the virus was metagenomically assembled. The middle ring refers to the sample site from which the corresponding sample was taken. All annotations of TerL in RefSeq and fumarole viruses were made with HHblits using parameters described in Methods.

We put forward two Caudoviricetes order-level clades containing 30 and 56 unique fumarole vOTUs respectively. These clades expand the diversity of dsDNA viruses and enable classification of three previously identified Caudoviricetes families which have not been assigned to orders previously. The first clade, which we call *Kilaueavirales*, contains the *Casjensviridae* and *Herelleviridae* families. The *Casjensviridae* family is made up of flagellotrophic phages which frequently infect Enterobacteriaceae and can be found in high abundances in wastewater (Chakraborty et al., 2024) while *Herelleviridae* viruses infect the Bacillota phylum of bacteria (Barylski et al., 2020). Clade 1 (*Kilaueavirales*) contains a family-level clade of 30 fumarole virus vOTUs (40 contigs) and 17 previously described RefSeq Caudoviricetes species which had not previously been assigned to viral families. These unassigned reference viruses have hosts such as *Xanthomonas*, *Burkholderia*, and *Enterobacter* (Black et al., 2025). This family-level clade of 30 fumarole vOTUs and 20 RefSeq viruses is monotypic with low support (support=60%) to the *Faecalibacterium* phage, *Eponavirus epona.* This family-level clade and the *Casjensviridae* family together form a well supported clade (support=100%) that is a sister group to family *Herelleviridae* (support=98.9%). Together, this resulting clade is Clade 1 (*Kilaueavirales*) (Supplemental Figure 02).

Clade 2 (*Pahoavirales*) contains 56 fumarole vOTUs, some of which make up clades which are sister groups to reference viruses (support=90.5%). One sister group within the *Pahoavirales* clade contains a fumarole virus with a monotypic relationship (support=89.1%) to RefSeq *Ralstonia*, *Agrobacterium*, and *Myxococcus* viruses. A sister group to this clade contains two sister subclades (support=95.8%). In one of these sister subclades, there are 24 fumarole virus vOTUs and viruses from the *Mesyanzhinovviridae* family are shallow-branching taxa close to tree tips. The second subclade (which is a sister group to the first subclade containing *Mesyanzhinovviridae)* is mostly made up of fumarole viruses (32 fumarole vOTUs), but also places two reference viruses (*Dinoroseobacter* and *Microbacterium* RefSeq phages) within the subclade (support=100%).The shallow branching relationships observed in the second sister subclade of *Pahoavirales* suggests recent divergence of closely related viruses without many close relatives (Supplemental Figure 02).

In addition to the 86 Caudoviricetes vOTUs which belong to two previously undescribed order-level clades, we observe other clades of Caudoviricetes fumarole viruses with some relation to previously identified viral families (Supplemental Figure 03). One fumarole vOTU is estimated to be within the same clade of Caudoviricetes incertae sedis viral family *Hodgkinviridae* (support=75.1%), which is a family of *Microbacterium* phages (Turner et al., 2025). We estimate 19 fumarole vOTUs (25 viral contigs) to be a sister group to the *Hodgkinviridae* family (support=93.7%). Six fumarole vOTUs are a sister group to the *Straboviridae* family (support=87.8%) (Turner et al., 2025). Five fumarole Caudoviricetes vOTUs are estimated to form a clade with the incertae sedis *Peduoviridae* family (support=85.9%). Two fumarole vOTUs are part of a clade with the *Zobellviridae* which is an incertae sedis Caudoviricetes family (support=96.5%). One fumarole vOTU forms a clade with the Caudoviricetes incertae sedis family *Saffermanviridae* (support=95.6%). The relationships of fumarole Caudoviricetes to numerous Caudoviricetes incertae sedis families provides evidence that the inclusion of these viruses in phylogenetic estimation expands the diversity of dsDNA tailed viruses.

#### Estimation of diverse Singelaviria and Varidnaviria

Viruses within the Singelaviria realm (class of Laserviricetes) and the Varidnaviria realm (class of Tectiliviricetes) are assigned to a single class category of ‘Laserviricetes|Tectiliviricetes’ by vConTACT3. Tectiliviricetes phylogeny was estimated with a maximum likelihood tree that was constructed using the packaging ATPase gene for six different fumarole Tectiliviricetes vOTUs (11 viral contigs). Viral species from all five genera within the *Tectiviridae* family were used as references. The resulting phylogeny shows six distinct fumarole Tectiliviricetes clades (Supplemental Figure 4). One vOTU is a sister group with weak support (support = 52.5%) to the Alphatectivirus genus, which contains only the two species of PRD1 and PRD4. Alphatectivirus is the only genus within the Tectiliviricetes class that is known to contain plasmid dependent phages (PDPs). This vOTU may be a taxon within Alphatectivirus, which is potentially worthy of further investigation when exploring the phylogenetic distribution of related ecological characteristics such as plasmid dependence (Bignaud et al., 2025). Two fumarole Tectiliviricetes vOTUs which are closely related and have almost identical packaging ATPase sequences to each other on the amino acid level are a sister group to the clade containing *Deltatectivirus* and *Epsilontectivirus* (support=93.8%).

An independent phylogenetic estimation with maximum likelihood was performed for Laserviricetes using the packaging ATPase gene. This phylogeny contains a single fumarole vOTU (Supplemental Figure 04) which is a sister taxon with strong support to the *Hukuchivirus* genus (support=100%). The *Hukuchivirus* genus is made up entirely of viruses infecting the thermophilic genus, *Thermus*. This vOTU could potentially be considered a member of a new genus within the family of *Matshushitaviridae*.

#### Microviridae form a poorly resolved clade with phiX174

A class-level maximum likelihood phylogeny of Malgrandaviricetes was built using the amino acid sequences of the major capsid protein (MCP) to classify the single fumarole *Microviridae* vOTU. The resulting phylogeny produced a poorly resolved clade, resulting in an apparent polytomy (support=100%) with Enterobacterial coliphage phiX174 (Supplemental Figure 05). These fumarole viruses and phiX174 are paraphyletic to the other genera of microviruses in the *Microviridae* family. The high genome conservation of phiX-like viruses (and more broadly, the *Bullavirinae* subfamily) (Kirchberger & Ochman, 2023) is potentially explanatory of an apparent polytomy with phiX174 with fumarole *Microviridae*, and suggests that phylogenetics is insufficient to capture the diversity of fumarole microviruses.

#### Comparative genomics of abundant Microviridae

A single vOTU of Microviridae was identified, with some genetic variation (subclustering) observed between phylogenetically identical Microviridae genomes of similar length (Figure 01). The relative abundance of Microviridae ranges from 22-82% across samples of varied microbial composition and biofilm phenotype (Figure 01). In almost half of biofilm samples, Microviridae makes up over half of the viral community, and is most abundant in moderately geothermal biofilm samples which have overall high viral richness and evenness. The high abundance of Microviridae in moderately geothermal biofilms is surprising because, to our knowledge, there has never been a terrestrial environment reported which displays high abundances of Microviridae. However, abundant Microviridae has been reported from viromic surveys of viruses in hydrothermal vent sediment, as well as benthic sediment (Cheng et al., 2022; Yoshida et al., 2018). Despite the more pronounced prevalence of Microviridae in moderately geothermal biofilms, mesothermal biofilms also display a higher abundance of Microviridae (>28% viral relative abundance) than both other terrestrial environments and mesothermal volcanic soil collected adjacent to geothermal features from the Kilauea caldera, suggesting Microviridae abundance is not simply predictable by measured temperature.

The low abundance of *Enterobacteria* in fumarole biofilms (Metagenome Assembled Genomes (MAGs) and Amplicon Sequence Variants (ASVs) suggest that these Microviridae have non-Enterobacterial hosts, and that the apparent polytomy of fumarole viruses with phiX-174 could be explained by low phylogenetic resolution rather than functional similarity between environmental *Microviridae* and phiX-174 (J. H. Saw et al., 2026). Our observation of viruses from the Bullavirinae subfamily in fumaroles is intriguing. While lab models of phiX-like viruses are heavily studied, the subfamily of Bullavirinae is considered extremely rare in nature and occupies a specialist niche as a lytic phage with limited capability for uptake of genomic content (Kirchberger et al., 2022).

Fumarole microviruses were characterized genomically with comparison of whole genomes (amino acid sequences) with RefSeq *Microviridae*. Network analysis with vConTACT3 demonstrates consistency in length of reference and fumarole *Microviridae* (phiX174 genome size is 5.386 Kb, fumarole *Microviridae* are 5.463 Kb), supporting that the potential for genome completion issues does not falsely inflate the degree of genetic difference between fumarole and reference *Microviridae*. Fumarole *Microviridae* are located within the same nested subcluster as viruses from the Bullavirinae subfamily, providing evidence of genetic similarity between fumarole viruses and Bullavirinae viruses beyond simply phylogenetic association. This overall genome conservation may be explained by limited genomic uptake capabilities of Microviridae, which occupies a specialist niche as a lytic phage, constraining genetic rearrangement and mosaicism (Kirchberger & Ochman, 2023). Despite this, fumarole and RefSeq Microviridae still vary widely with pairwise similarity as low as 40-60% between viruses (Figure 03). In addition to low overall genome similarity, there is evidence of syntenic changes between fumarole Microviridae, with differences from RefSeq Bullavirinae species (Supplemental Figure 06).

**Figure 03.**
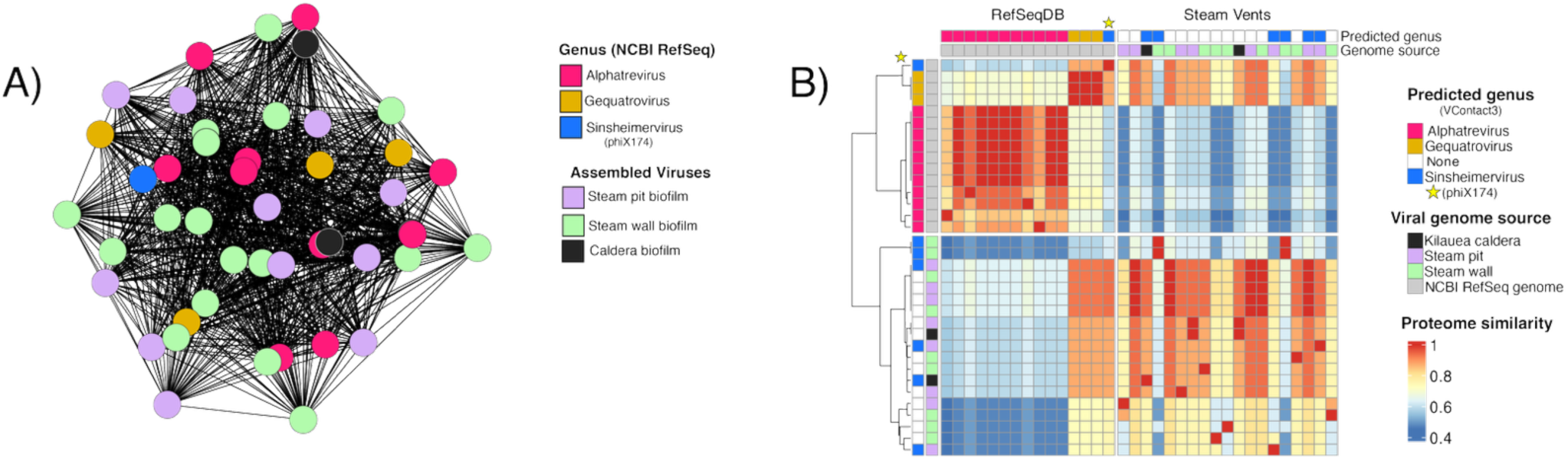
**A)** Subcluster of viral genomes within the family Microviridae. Network constructed using vConTACT3 and visualized in Cytoscape. Reference nodes are colored by genus (RefSeq and vConTACT3 predictions), while fumarole viruses are colored by sample site. Edges represent gene-sharing strength. The blue node (phiX174) forms a weakly resolved clade with fumarole viruses in the maximum-likelihood phylogeny based on major capsid protein amino acid sequences. **B)** Pairwise genome comparisons of RefSeq and fumarole Microviridae using local alignment (MMseqs2). The top row and left column indicate predicted genus assignments (vConTACT3) for fumarole and RefSeq viruses. The top-left quadrant shows RefSeq–RefSeq comparisons; the top-right shows fumarole–RefSeq comparisons (RefSeq colored by genus). The second row (genome source) is colored by fumarole sample site. The star denotes comparisons involving the phiX174 RefSeq genome.

The low pairwise similarities and syntenic rearrangement we observe between fumarole *Microviridae* is highly unexpected. While innovations do occur in the *Microviridae* genome (gene overprinting, recombination) the overall high syntenic conservation of the *Microviridae* family is thought to reflect strong evolutionary constraints for transcriptional timing of *Microviridae* viruses (Kirchberger & Ochman, 2023).

### Biogeography and diversity of vOTUs

#### Most vOTUs are shared between biofilms

All viral contigs were mapped to samples and normalized by calculating reads-per-kilobase-per-million (RPKM). Contigs were summed for each vOTU and VC for alpha-diversity analysis and visualization respectively (Roux et al., 2019; ter Horst et al., 2021). 99.7% of vOTUs (n=382) are shared between biofilm samples (all but one vOTU). The observation that most vOTUs are not unique to individual biofilms or sample sites is consistent with predictions by the Bank model (Breitbart & Rohwer, 2005). However, evidence of viral dispersal is unexpected within the physicochemical context of spatially separated and geothermally-active fumaroles that give way to harsh conditions (J. H. Saw et al., 2026). From hyperthermophilic hydrothermal vents to moderately geothermal hot springs, both microbes and viruses are found to be overwhelmingly endemic (Jarett et al., 2020; Langwig et al., 2025; Salgado et al., 2025). Both Hawaiian fumarole-hosted microbial communities (Prescott et al., 2022) and viral communities show evidence of selective pressure associated with the geothermal activity. Within Big Ell samples, there are few shared vOTUs between the rock-hosted thermal biofilms and volcanic soil, collected directly adjacent to the cave wall. No vOTU which is found in over 5% abundance within thermal biofilms has an abundance over 1% in any non-thermal volcanic soil sample (Figure 01). The vOTU communities of thermal biofilms is suggestive of selection, rather than random vOTU dispersal (Liang et al., 2019).

Yet, while microbial communities of fumaroles form “islands” with distinct communities between biofilms of different thermal features (Prescott et al., 2022), shared vOTUs are observed in unexpectedly high abundances across samples. 9.5% of vOTUs (n=36) are shared between samples with individual population relative abundance over 1% (Figure 01). 5% of identified vOTUs (n=19) are shared between geothermal features and sites in abundances over 2%. The abundance of shared vOTUs between fumaroles is inconsistent with descriptions of distance decay observed in other physicochemically diverse geothermal systems, which are harsh to varied extents. For hot spring and hydrothermal vent systems, viral community dissimilarity increases with geographic distance, and fewer shared vOTUs are observed between distant geothermal fields (regions with localized and relatively high heat flow). Though the seed-bank model predicts that most of the shared vOTUs will be found in <2% relative abundance within a given community at any point in time, (Hevroni et al., 2020) we observe individual fumarole-hosted viral populations that are non-dominant and shared between samples can make up to 10% of a given biofilm’s viral relative abundance, representing a significant variation from the Bank model.

Perhaps as a consequence of dispersal limitation, specific host virus pairs are often observed in high abundances within geothermal systems (Guajardo-Leiva et al., 2021a; Salgado et al., 2025; R. Wu et al., 2023). This has been described as particularly prevalent in phototrophic thermal biofilms of hot springs, where matrix immobilization of microbes and their viruses result in dispersal limitations and facilitates tightly coupled host-virus pairs (Jarett et al., 2020). While it is difficult to determine microbial and viral mobility metagenomically, CRISPR-based analysis provides evidence of coevolutionary relationships between microbes and their viruses in line with the Red Queen hypothesis (Common et al., 2019; Koonin & Wolf, 2015; R. Wu et al., 2023). For the 10% of recovered fumarole vOTUs which were host matched (n=39), specialized host-virus relationships were determined by the presence of more than three unique spacer matches between a MAG and a vOTU. Out of the 21% of medium to high quality MAGs (n=74) which were successfully host matched, 11 CRISPR arrays showed evidence of microbial coevolution with vOTUs (J. H. Saw et al., 2026). The most spacer matches (n=61) within a single array comes from a Rhodomicrobiaceae MAG from an unclassified genus. MAGs from the Chloroflexota families Roseiflexaceae (n=35) and JAJPGT01 (n=34) also provide CRISPR-based evidence of an antagonistic coevolutionary relationship between specific vOTUs and their hosts.

Antagonistic coevolution described by ‘Kill-the-Winner’ is inherently density dependent (Hu et al., 2025; Salgado et al., 2025). Despite this, not a single of the 14 vOTUs which is identified as part of a specialized host-virus pair can be found in viral relative abundance over 1%. The associated host phyla abundance also does not correspond to linked vOTU abundance (Supplemental Material S6) (Prescott et al., 2022; J. H. Saw et al., 2026). This unexpected low abundance is observed across microbial taxa with specialized pairing to vOTUs. Cyanobacterial genus *Chlorogloeopsis* is highly abundant in steam wall biofilms, and demonstrates a multi-spacer relationship with a highly novel paired vOTU which is below relative abundance of 1% in all samples except for steam wall biofilm PDB5 (9.24% relative abundance) non-thermal volcanic soil sample HW-02, where it is found in 15.3% relative abundance. These samples are not in close proximity to each other and have distinct microbial community composition suggesting that Hawaiian fumarole viruses and their hosts are not necessarily endemic, even though these shared viruses participate in specialized host-virus pairings (Audrey et al., 2023). These spacers are not always located in the same array. We observe spacers paired to a vOTU can be distributed in multiple arrays across the host genome. This phenomena could be evidence of repeated exposure to a particular vOTU, or be a result of multiple different CRISPR systems conferring viral resistance (Shmakov et al., 2018). The distribution of vOTU resistance across two arrays is observed for three Chloroflexota MAGs from the Chloroflexia and UBA6077 classes, which contain various Type I and Type III systems (Supplemental Material S7). These arrays are not necessarily genomically clustered, with up to ∼61 kb observed between these array pairs.

While the majority of samples do not have high abundances of identified tightly-coupled virus-host pairs, high relative abundance is observed for two Caudoviricetes vOTUs which are both prophages belonging to MAGs of *Gloeobacter kilaueensis*. This cyanobacteria is abundant only in steam pit samples, which is observed for these vOTUs as well. High abundance of Gloeobacter prophages is only observed as expected in samples dominated by *Gloeobacter kilaueensis*. Summed vOTUs occur in relative abundances that range from 17-51% in all but one of the biofilm samples with high abundance of microbial *Gloeobacter kilaueensis*. These two prophage vOTUs follow the directly proportional abundance trends predicted by the “Piggyback-the-Winner” hypothesis which describes a microbial community in which co-abundant vOTUs favor a switch to lysogeny as a viral life cycle strategy, and accordingly, abundant viral populations are identified as prophages (Hu et al., 2025). The presence of Gloeobacter prophage vOTUs is in line with this theory. One of these vOTUs is present in double the abundance of the other vOTU across all samples (Supplemental Material S6). Unlike many of the other vOTUs with high dispersal inferred (abundance in samples doesn’t correspond to either hosts or microbial taxa) these two vOTUs appear restricted in dispersal to be consistent with host.

#### Viral alpha-diversity

Chao1 (species richness) and Pielou (species evenness) index values were plotted against each other for each sample (Figure 04). Moderately geothermal biofilms cluster together, regardless of location or microbial distribution within the sample. Relative to other samples, biofilms which are collected from moderately geothermal fumaroles (40-60℃) have high richness and evenness. The 17 thermal biofilm samples show a strong positive correlation between vOTU richness and evenness. On the plot of richness to evenness, non-thermal volcanic soil diversity is clearly partitioned from the fumarolic active biofilms, displaying low vOTU richness and high evenness. The low richness is striking when compared to biofilm diversity, because soil has been described as a system that generally has high viral richness and evenness comparative to the viral communities of other systems such as marine and groundwater (Roux & Emerson, 2022; ter Horst et al., 2021).

**Figure 04.**
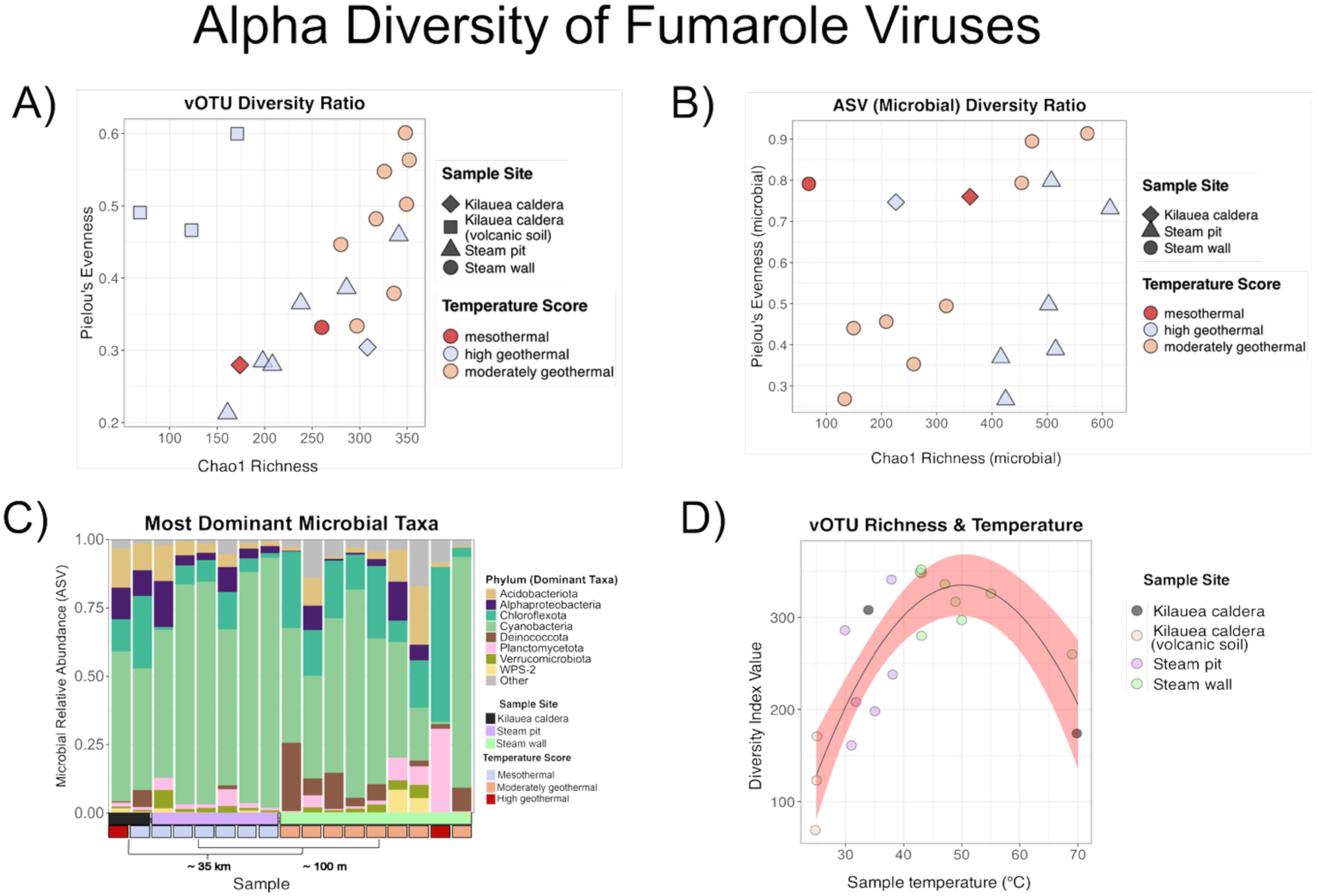
**A)** Richness (Chao1) and evenness (Pielou) of fumarole vOTUs calculated from RPKM-based relative abundances (Supplemental Material S6). Each point represents a sample. Shape indicates sample site, and color indicates temperature at time of collection. **B)** Richness and evenness of microbial ASV relative abundances. Each point represents a sample; color and shape denote sample characteristics as described in (A). **C)** Relative abundance of the most abundant bacterial taxa across samples based on ASVs. The top five phyla per sample are shown and used for visualization. The second row indicates sample site, and the bottom row indicates temperature at collection. The phylogram shows spatial relationships among sites: the steam pit and steam wall are ∼100 m apart, while the Kīlauea caldera cave is ∼35 km from both. **D)** Sample temperature versus vOTU richness (Chao1). Each point represents a sample, colored by site. vOTU richness follows a temperature-dependent pattern modeled by a polynomial (quadratic) regression. Richness increases with temperature up to ∼50°C and declines thereafter.

Richness and evenness plots exhibit a clear partitioning of diversity (high richness/evenness in moderately geothermal fumaroles) that suggests diverse viral communities may be characteristic of active fumaroles. The biofilms which display this high species richness and evenness are dominated by different microbial taxa, suggesting that microbial taxonomy alone is insufficient to explain viral diversity (Figure 04). Microbial diversity indexes of biofilms were assessed using ASVs (Prescott et al., 2022; J. H. Saw et al., 2026). Unlike with viruses, there is no visually evident clustering of temperature-associated diversity (Figure 04). The decoupling of microbial and viral diversity is observed across samples of varied temperature ranges and microbial compositions. The samples with high viral richness and evenness are composed of diverse phyla (Figure 04). Interestingly, these dominant phyla all are known to occupy varied niches and display varied metabolic traits, suggesting microbial composition alone is insufficient to be the primary explanatory driver of viral diversity within these biofilms (Prescott et al., 2022).

When the relationship between richness and temperature was assessed, a quadratic model was found to be the best fit to explain this relationship. While the 4-parameter cubic model had the lowest residual sum of squares (RSS) value (RSS=36846.4), the quadratic model had the lowest BICc value (BICc=161.8) indicating that the quadratic model is the best fit for describing the relationship between temperature and viral richness for non-thermal volcanic soil and fumarolic biofilms, ranging in temperature from mesophilic to highly geothermal (>70°C) (Prescott et al., 2022; J. H. Saw et al., 2026). These tests support the significance of a non-linear (quadratic) relationship between temperature and viral richness in the Hawaiian fumarole system, with the highest vOTU richness observed at ∼50°C. The extent to which microbial life adheres to Humboldt’s theory of a temperature-diversity gradient has only been explored in a handful of geothermal environments, with hot springs making up the majority of geothermal systems studied for this (Sharp et al., 2014). In some moderately geothermal circumneutral hot springs, the highest species richness of bacteria has been observed at 50°C, with increasing temperatures associated with decreased species richness (Castelán-Sánchez et al., 2020). While the bacterial diversity of fumarolic biofilms observed does not clearly follow this trend, viral diversity demonstrates a surprisingly consistent temperature-diversity gradient which is independent of microbial diversity (Prescott et al., 2022; Sharp et al., 2014). This indicates that moderately geothermal features of fumaroles are potential biological hot spots for viruses, with high biodiversity of microbial/viral life which is observed across different geothermal systems at similar temperatures.

The R-squared value (R-squared=0.7292) indicates temperature explains ∼73% of variance in viral richness, and is determined to be significant with the p-value of the F-statistic (p=1.619e-05) (Figure 04). Normality of residuals was supported by QQ and density plots, and the Shapiro-Wilk test confirms that the quadratic model meets normality assumptions (W=0.94514, p-value=0.2992) (Supplemental Figure 07). Though vOTU evenness varies greatly between samples (from Pielou=0.21-0.6), temperature cannot explain the great variation in viral population evenness (Supplemental Figure 07). While there is a significant relationship between vOTU richness and temperature (consistent with the cluster of high viral richness and evenness for moderately geothermal biofilm samples) the evenness of these samples cannot be described as temperature dependent, or consistent with any specific phylogenetic distribution, indicating complex biogeographic factors may explain viral diversity of biofilms.

### CRISPR defends fumarole bacteria against a breadth of unknown elements

CRISPR spacer targets (n=302) were identified for arrays from 13 different microbial phyla (Supplemental Material S7). Matches are found for 20% of all fumarole-assembled MAGs (n=74) (J. H. Saw et al., 2026). The majority of spacer matches (n=120) are for arrays that come from Chloroflexota MAGs (classes:UBA6077, Ktedonobacteria, and Anaerolineae). Despite this, only 70% of the Chloroflexota targets identified are vOTUs (n=9). Additionally, Chloroflexota spacer matches were identified for 1 plasmid (0.65% of all spacer matches) and 33 unknown elements. Beyond the Chloroflexota phylum, spacer targets are not limited to viruses either. Even though the majority of matches identified were to vOTUs targeted by spacers (74.2% of spacer matches) viruses were only 44% of the matched target elements.

These unknown elements represent possible other miscellaneous mobile genetic elements (conjugative/integrative elements, transposons, insertion sequences) (Tokuda & Shintani, 2024) or chromosomally integrated elements (Shmakov et al., 2017). Even though the majority of spacer targets are non-viral, these unknown element matches make up only 20.86% of total spacer matches. This suggests that the microbial populations of fumaroles utilize CRISPR to defend themselves against a wide breadth of unknown elements, and that CRISPR may play a minimal role in any sustained antagonistic coevolution occurring between non-viral elements and microbes in the fumarole biofilms. (Martynov et al., 2017). One exception to this, however, is multiple spacer matches (n=8) between an unknown element and a Chloroflexota MAG (class UBA6077). Seven of these matches to the unknown element are part of a Type IA CRISPR system, while one spacer match is contained within a different array that is part of a Type IE CRISPR system.

### Identification of putative AMGs and accessory viral genes

#### AMG mediated adaptation to heavy metal toxicity and resource limitation

Secondary curation of baseline DRAM-v AMG summary hits (212 genes) was performed to eliminate viral life cycle and host-associated genes from AMG candidates. This yielded 72 putative AMGs and 70 accessory (moron) genes associated with defense against mobile genetic elements (Brüssow et al., 2004). The majority of AMG candidates and accessory defense genes we identify are from viruses in fumarole biofilms - only one accessory defense gene and one AMG candidate is observed in viruses assembled from non-thermal soil. Overall, 7% of vOTUs (n=27) identified from fumarole metagenomes contain AMG candidates. These low AMG frequencies have been observed in other geothermal systems such as hydrothermal vents (Langwig et al., 2025). The majority (74%) of AMG-carrying vOTUs are classified as Caudoviricetes, and the remaining vOTUs with AMG candidates cannot be assigned to realms (Figure 05). Five vOTUs contain AMG-candidates related to energy metabolism (boosting ATP-generation). Three of these vOTUs encode NifU-like IscA (scaffold and iron carrier) which is a gene involved in Fe-S cluster biosynthesis. One vOTU contains AMG candidate gene lactate dehydrogenase (reversible enzyme that converts pyruvate to lactate to generate NAD+ for glycolysis, or converts pyruvate to lactate, feeding the TCA cycle). Another vOTU contains putative AMG succinate dehydrogenase cytochrome b558, which, to our knowledge, has only been described as an AMG in giant virus family Mimiviridae (Blanc-Mathieu et al., 2021; Lamb et al., 2024).

**Figure 05.**
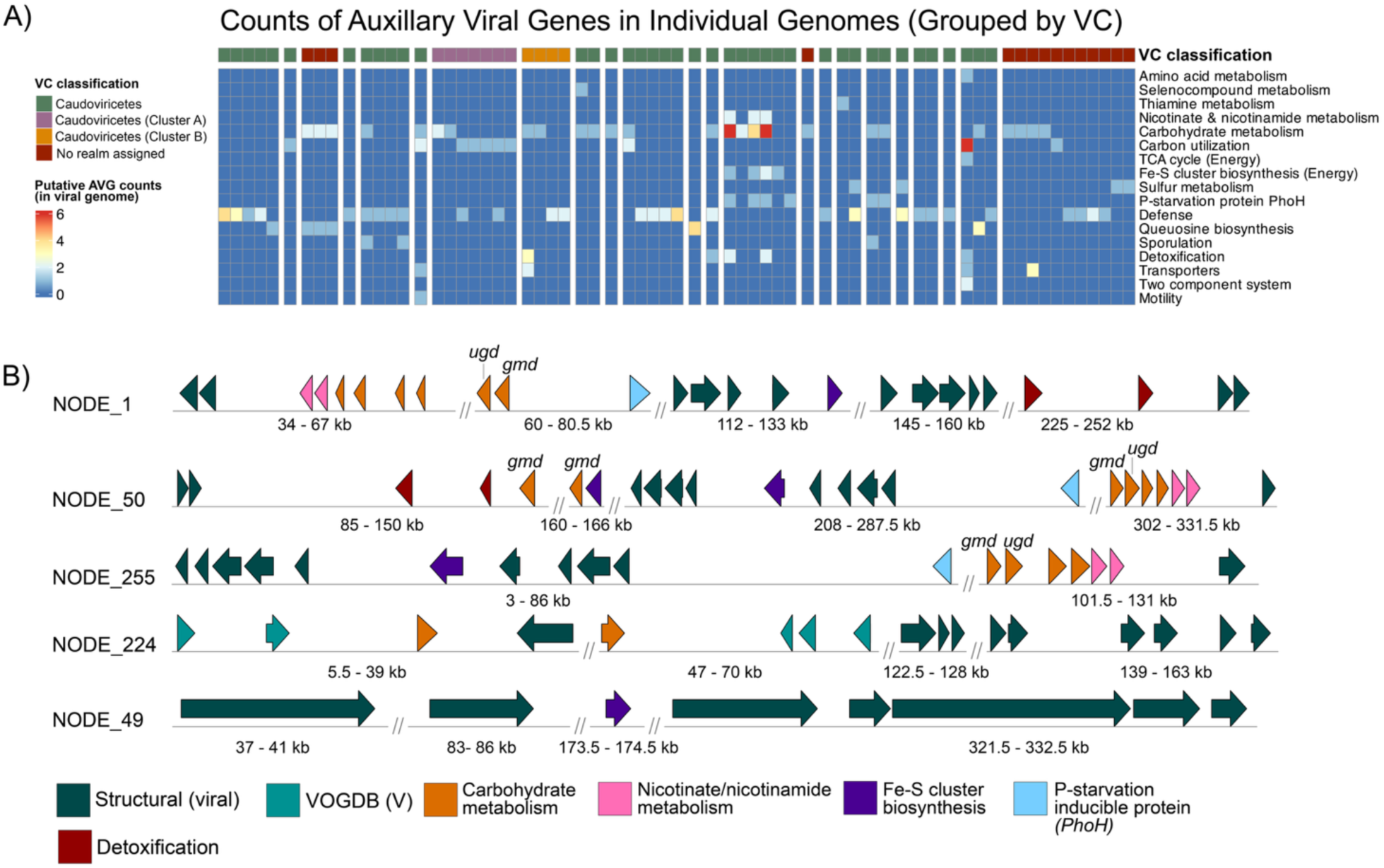
**A)** Counts of viral accessory genes in individual viral genomes. Columns represent viral metabolic gene counts per contig. The top row indicates vConTACT3 viral cluster (VC) assignments, with spacing grouping viruses within each VC; singletons are shown as individual VCs. Heatmap colors denote gene counts as shown in the legend. **B)** Gene diagrams of viral metabolic gene distribution across five recovered genomes from vOTU 158, enriched in putative AMGs. Genomes were inspected for flanking viral genes and AMG annotation coverage/identity. Arrows denote genes, and colors indicate AMG categories. The full curated AMG list is provided in Supplemental Material S8.

The majority of putative AMGs identified in fumarole viruses encode transporter-related gene products (Supplemental Material S8). Transporter types include: divalent metal (DMT) transporters, RND efflux pumps, ABC-transporters, and Major facilitator superfamily (MFS) transporters. Ten fumarole vOTUs contain AMG candidates that can be linked to cellular detoxification from heavy metals, such as metallothioneins, Zn2+/Cd2+ exporting ATPases, and heavy metal efflux proteins including transporter TolC. The TolC transporter functions with efflux of transition metals, including first row transition metals which are present in basalt (Farias et al., 2015). The transporters observed in fumarole viruses have also been described in AMG candidates from viruses in hydrothermal vents and heavy metal enriched soils with similar elemental composition to basaltic surfaces described in the ERZ volcanic system (Prescott et al., 2022; Sun et al., 2023; Thomas et al., 2021). Other fumarole virus transporters are protective proteins (such as Fe-Mn superoxide dismutase) that neutralize reactive oxygen species (ROS) which can also be generated by metal induced cellular stress. Fumarolic non-transporter AMG candidates (such as rubrerythrin) also neutralize ROS, which usually confers host survival in microaerophilic or anoxic environments (Cardenas et al., 2016).

Other AMG candidates relate to nutrient deficiency. One vOTU contains sporulation maturation protein CgeB which is sometimes observed in Bacillus subtilis prophages where it is part of the Cge operon which modifies the surface properties of the spore (Shuster et al., 2019). One putative vOTU contains a carbohydrate ABC transporter permease (Trehalose transport system permease protein SugB) which is involved in trehalose recycling within the cell (Pohane et al., 2021). One fumarole vOTU (prophage of Gloeobacter kilaueensis) encodes a gene highly similar to polyhydroxybutyrate depolymerase (phaZ). Polyhydroxybutyrate depolymerase is an intracellular enzyme which breaks down polyhydroxyalkanoates (PHAs) or Polyhydroxybutyrate (PHB) biopolymers which can be used as carbon sources. Eight unique vOTUs contain AMG candidates related to carbon utilization. Two Caudoviricetes vOTUs contain five different carbon utilization genes that could possibly be involved in PHA biosynthetic pathways, converting short-chain carboxylic acids (such as acetate, propionate,lactate, and butyrate) into PHA (Yao et al., 2025). Alternatively, these five genes could be involved in beta oxidation of fatty acids, which have been described in the giant virus family Mimiviridae (Brahim Belhaouari et al., 2022). The presence of these five AMG candidates in Caudoviricetes as observed in two fumarole Caudoviricetes vOTUs is unusual because Caudoviricetes hosts are prokaryotic while Mimiviridae hosts are eukaryotic (Duda & Teschke, 2019). Another putative AMG contained within one of these Caudoviricetes vOTUs is gene 2-methylcitrate dehydratase (prpD) which catabolizes the third step of the methylcitrate cycle for propionate catabolism, and has been described as an AMG in hydrothermal vents (Zheng et al., 2021). Host linking with experimental verification of transcriptional activity and gene product expression is necessary to ascertain if these putative AMGs are involved in beta oxidation of fatty acids and lipid catabolism pathways, or PHA biosynthesis pathways.

#### Accessory vOTU genes may confer resistance to non-self viruses

In addition to AMG candidates, 32 vOTUs (8% of fumarole derived vOTUs) contain virally encoded defense genes, which can potentially provide antiviral defense for the microbe (increasing host fitness), or facilitate superinfection exclusion against other viruses. 78% of the defense-containing vOTUs (n=25) were classified as Caudoviricetes, and the remaining defense-containing vOTUs (n=7) could not be assigned to a realm classification. Queosine biosynthesis genes (7-Deazaguanine modification) and (cytosine-5)-methyltransferase (dcm) were initially flagged by DRAM-v as metabolic AMG candidates. However, they were manually reclassified as defense-related because that is more likely reflective of functional role (Hutinet et al., 2019; Wang et al., 2024).

Fumarole viruses contain a range of defense and anti-defense related genes: anti-CRISPR, anti-Thoeris, and 7-Deazaguanine modification genes were annotated by DRAM-v. Virally encoded toxin/antitoxin (TA) genes from Type I-IV systems were observed in ∼5% of total fumarole vOTUs (n=18) (Supplemental Material S8). For most of the vOTUs (n=10) containing TA systems, both toxin and antitoxin genes of the same system type are positioned with no more than five genes between components, suggesting that at least some fumarole viruses encode functional TA systems. Two vOTUs contain CRISPR-associated proteins. One vOTU (vOTU 189) does not have an identified host and encodes a minimal Type I-C CRISPR-Cas Cascade system, which contains only three unique Cas proteins: Cas5d, Cas8c, and Cas7. This minimal Cascade encodes an RNA-guided multi-subunit CRISPR-Cas surveillance complex which targets foreign DNA for destruction (O’Brien et al., 2022).

A different vOTU (vOTU 212) contains a minimal Type IV system (genes Cas6, Csf3, and Csf2) and an unassociated CRISPR array. Type IV systems are primarily encoded by plasmids and more occasionally observed on prophages (Moya-Beltrán et al., 2021). Type IV CRISPR systems are enigmatic in mechanism, and usually lack a CRISPR array. They are hypothesized to modify existing CRISPR-Cas activity of hosts which already contain CRISPR systems (Koonin & Makarova, 2017). Accordingly, vOTU 212 is targeted by spacers within a Chloroflexota (class UBA6077) MAG which contains both Type IA and Type IE systems (J. H. Saw et al., 2026). In addition to the Type IV effector proteins, vOTU 212 encodes a CRISPR array. The maintenance of both an array and Type IV system (which characteristically, lack associated CRISPR arrays) (Pinilla-Redondo et al., 2020) is suggestive of inter-viral competition which has resulted in the use of CRISPR as a method for viruses to prevent superinfection exclusion (Muscatt et al., 2022).

## Discussion

### Fumaroles contain novel and diversifying viruses

Our approach to viral identification that utilizes sensitive, profile-based methods (Steinegger et al., 2019) for annotation of structural viral gene components reveals novel viruses from fumarole biofilms and volcanic soil. Our approach identifies viruses using amino acid sequences encoding viral structure components (capsid-associated and packaging genes) which are highly divergent (Söding et al., 2005). Our recovery of phylogenetically novel viruses from biofilm metagenomes and our subsequent careful manual verification supports our workflow which utilizes remote homology protein prediction for identifying structural virion components. This can, in turn, be used for identification of viruses in a high-throughput processing pipeline with existing tools (Felipe Benites et al., 2024) without extensive false positives (Supplemental Material S4).

Our approach yields recovery of highly novel viruses (Bolduc et al., 2025) which we find to be sometimes highly abundant relative to other viral populations (Figure 01). This is predictive of either functional significance for these novel and abundant viruses, or represents significant population dormancy (Liang et al., 2019). Because viruses do not all share a single gene that is common to all species (Harris & Hill, 2021), there is not a way to confidently place all viruses of microbes on a single phylogenetic tree of life. Thus, it is not possible to phylogenetically estimate viruses which cannot be classified into an existing realm (Koonin & Wolf, 2012). Our assessment of viral diversification for these ‘no-realm’ viruses relies on gene-sharing network analysis, which provides a method for comparison of genomic distance between viruses which cannot be otherwise taxonomically placed. While uncertainty in phage genome completion (vOTUs of variable length within a cluster) and chimeric or mosaic assemblies can constrain comparisons of viral genome similarity, our utilization of NetworkAnalyzer metric of ‘Closeness Centrality’ (Su et al., 2014) minimizes the effects of incomplete viruses producing erroneous inferences of genomic distance (Bolduc et al., 2017). Our observation of subclusters for these highly novel viruses, which are not placed into a realm by vConTACT3 and do not exhibit gene sharing with previously described viruses, is evidence that there may be genomic diversification between these novel viruses. Yet, conclusions about associated viral speciation for these highly novel populations are limited by our inability for phylogenetic estimation without realm classification.

The majority of recovered viruses are from the Caudoviricetes class. This could possibly be a result of detection facilitated by high representation of Caudoviricetes genomes within databases of known viruses (Weinheimer et al., 2025). As a result, our conclusions about evolutionary patterns in other viral classes are limited by the low genomic diversity of recovered fumarole vOTUs from the Laserviricetes, Tectiliviricetes, and Malgrandaviricetes classes. Even though Caudoviricetes are the most characterized phage class, our identification of fumarole viruses significantly expands the diversity of the class. Out of the 179 Caudoviricetes vOTUs included in phylogenetic estimation, 86 fumarole vOTUs belong to two previously undescribed order-level clades within the Caudoviricetes class (Supplemental Figure 02). The 93 other fumarole Caudoviricetes vOTUs have varied levels of relation to previously described viral families. Phylogenetic estimation exhibits that many of the fumarole-derived Caudoviricetes vOTUs are related to families which are primarily incertae sedis within the Caudoviricetes class such as *Hodgkinviridae*, *Peduoviridae*, *Zobellviridae*, and *Saffermanviridae* (Supplemental Figure 03). This suggests that the inclusion of novel viruses may help resolve the phylogenetic positions of problematic taxa within the Caudoviricetes class. Our observation of distinct Caudoviricetes fumarole vOTUs which have close phylogenetic relationships (Figure 02) suggests not only genomic diversification, but also speciation, could be occurring between fumarole Caudoviricetes populations.

### Abundant Microviridae may be a signature of geothermal environments

Microviridae are known to be diverse and globally distributed across a variety of environments (sediment, marine, animal, insect, and humans) (Kirchberger et al., 2022; Kirchberger & Ochman, 2023; Roux et al., 2012). Despite this, there has never been a description of a terrestrial environment in which Microviridae is the most abundant viral population, or otherwise represents a substantial portion of viral abundance in terrestrial or marine (oceanic zone) environments. Benchmarking with mock communities representative of open ocean marine and freshwater environments are consistent with observations that ssDNA viruses make up a very small portion (<5% total DNA viruses) of total viral content, and are much less abundant than dsDNA viruses (Kirchberger et al., 2022; Kirchberger & Ochman, 2020). The other environment in which Microviridae has been described as highly abundant is in deep ocean hydrothermal vents sediments (Castelán-Sánchez et al., 2019; Cheng et al., 2022; He et al., 2017; Y. Wu et al., 2024). It is possible that viruses belonging to Microviridae can be adapted to thermal conditions as an ssDNA virus. Phage phiX-174 has been demonstrated to be most stable at 37 - 44°C and is predicted to be favored over dsDNA and RNA phages under thermal stress because of the increased access that ssDNA viruses have to mutational diversity (Greenrod et al., 2025). Alternatively, it is possible that Microviridae has the capability to circularize which would enable abundance at higher temperatures (Laanto et al., 2017). However, the ERZ fumaroles with highest Microviridae abundances (50-70°C) are over three times cooler than sediment of Southwest Indian Ocean hydrothermal vents (temperatures as high as 400°C), where Microviridae has been identified as the most abundant virus (Cheng et al., 2022).

Out of ERZ fumarole viral communities, moderately geothermal and high geothermal biofilms are described as having the highest Microviridae abundances out of all samples. However, mesothermal fumarole biofilms are observed to have high Microviridae abundance (22-28%) which is higher than described in other terrestrial and marine environments (Figure 01). This may be explained by the high abundance of Microviridae in mesothermal fumaroles experiencing rapid fluctuations in fumarole temperature. Mesothermal fumaroles which harbor high Microviridae abundance may reflect a lag in decreasing abundance of ssDNA microvirus abundance as the biofilm communities adapt to lower temperatures. Volcanic activity of the ERZ varies widely across small spatial scales and fluctuates rapidly within short time spans (minute to minute), making it possible that the steam pit mesothermal fumaroles are dying geothermal features, while the ∼100 m adjacent steam wall is more geothermally active. Many fumaroles are ephemeral and experience more variation in activity as they near the end of their geothermal life span (Madonia, 2020).

Viromics is known to overestimate the abundance of ssDNA viruses, and the perception of high abundance is often a sequencing artifact. Many viromics extraction approaches may utilize Adaptase prior to linker amplification and produce a quantitative bias, leading to erroneous predictions of microvirus abundance (Roux et al., 2016). Alternatively, many viromics DNA preparation methods for low biomass viromics may destroy Microviridae DNA (Gregory et al., 2019; Y. Wu et al., 2024). Our approach utilizes bulk metagenomics prior to sequencing, which may explain the abundance of microviruses we observe. We hypothesize that microvirus diversity is underpredicted in geothermal environments, and temperature may be a selective pressure which favors ssDNA viruses (Roux et al., 2016). Studies of other terrestrial geothermal systems have not yielded similar observations of dominant Microviridae. However, some of these other systems (such as terrestrial hot springs) are often explored in the context of low biomass sediment that requires amplification or alternative methods of extraction which may destroy microviruses (Gregory et al., 2019). Alternatively, these geothermal systems are investigated with methods of viromics that destroy ssDNA viral representation in the sample (Roux et al., 2012). Fumarole biofilms may not be the only terrestrial system whose viral community contains dominant *Microviridae*, but the high fumarole biofilm biomass which is utilized for DNA extraction may reveal this characteristic of thermal viral communities which is otherwise obscured by methodological decisions which are made when investigating the microbial and viral diversity of other terrestrial thermal systems.

Fumarole *Microviridae* are highly novel with pairwise genome similarities as low as 40-60% (Figure 03) as well as syntenic rearrangement between viruses within the fumarole biofilms and compared to reference *Microviridae* (Supplemental Figure 06). However, there is a need for comparative genomic approaches when analyzing highly conserved ssDNA viruses to make conclusions about the degree of novelty introduced with fumarole *Microviridae*. Network analysis with vConTACT3 demonstrates multiple *Microviridae* form a single subcluster which is unexpected because clusters usually represent genus level rank (Bolduc et al., 2017, 2025). This suggests that environmental *Microviridae* are highly conserved and/or significantly under-described.

### Viruses are widely shared between geothermal features of ERZ fumaroles

There is evidence that viral dispersal has consequences for fumarole microbial evolution. Antagonistic coevolutionary relationships between hosts and viruses are often described through a ‘Kill-the-Winner’ dynamic, where both host and virus engaging in an active arms race are abundant relative to other viral and microbial populations. Relatedly, viral dispersal is thought to be host-dependent (Langwig et al., 2025) and associated with microbial abundance (Salgado et al., 2025). Despite these expectations, we observe many of the shared vOTUs are only found in low abundances within samples. For microbes that appear to utilize CRISPR as a defense mechanism and have identified arrays, vOTUs were determined to be part of a ‘specialized’ host-virus pair if there were more than three spacers linking a vOTU to a single microbial host (indicative of antagonistic coevolution). The 14 vOTUs which have specialized pairings with microbial hosts are not found in viral relative abundance over 1% within any sample. This suggests that low abundance viruses and hosts are still antagonistically coevolving and dispersal. Host-dependence and abundance are insufficient factors to predict limited dispersal of viruses in fumaroles. However, it should be noted that many microbial taxa do not rely on CRISPR for viral defense, and viral novelty constrains capability for host identification of 90% of identified fumarole vOTUs (Tan et al., 2024). Yet, even when simply searching for patterns of concurrent variation between abundances of unmatched vOTUs and hosts, no high abundance concurrent variation that would suggest an expected ‘Kill-the-Winner’ trend was observed (Supplemental Material S6).

Most vOTUs being shared between fumarole biofilms is highly unexpected. For hot springs and hydrothermal vents, individual vents and pools are biological “islands” with distinct communities between features (Castelán-Sánchez et al., 2019). Microbial communities of Hawaiian fumarole vents exhibit distinct communities as well (Prescott et al., 2022). In hot spring and hydrothermal vent systems, viruses display a distance decay effect with fewer shared viral populations observed with increasing distance between features (Jarett et al., 2020). The majority of viruses being shared between samples is even more unexpected when the context of phototrophic biofilm matrixes is considered (Castelán-Sánchez et al., 2020; Jarett et al., 2020). Compared to other mediums of geothermal systems (sediment or water), geothermal biofilms with strict spatial stratification are expected to have even less dispersal, with deterministic selection occurring on viruses within the microscale of centimeters within the biofilm (Guajardo-Leiva et al., 2021a). This is particularly remarkable considering the context of this comparison for fumarole biofilms to hot spring phototrophic biofilms with similar biogeochemical cycling processes and organismal characteristics (Guajardo-Leiva et al., 2021b; Salgado et al., 2025).

The high viral dispersal that is suggested by many shared vOTUs could be explained by unique dispersal mechanisms of fumaroles that diverge from the methods of dispersal seen in hot spring and hydrothermal vent systems. Steam (groundwater heated by magma) may be a mechanism for viral dispersal (Wall et al., 2015). Steam may enable biofilms to act as “extremophile collectors” which retain viruses within biofilms (Prescott et al., 2022; Wall et al., 2015). This suggests Hawaiian fumaroles in the ERZ have unique ecological and complex biogeographic characteristics with high viral dispersal informing these communities. By elucidating the metabolic and phylogenetic diversity of fumarole viruses, we have identified ERZ fumaroles on the Big Island of Hawaìi as a promising and unique early Earth analog environment to investigate viral ecology.

### *Gloeobacter* prophage may increase host fitness

The steam pit sample site is dominated by *Gloeobacter*, despite the close proximity and presence of other oxygenic phototrophic Cyanobacteria (J. H. Saw et al., 2026) which are faster-growing and photosynthetically efficient compared to *Gloeobacter* species which contain reduced photosystems (Grettenberger, 2021). Two vOTUs (Supplemental Figure 08) identified as *Gloeobacter kilaueensis* prophages by global alignment to *Gloeobacter* MAGs (J. H. Saw et al., 2026; J. H. W. Saw et al., 2013) are highly abundant and co-vary with *G.kilaueensis* in abundance for all but one of the biofilm samples dominated by *G.kilaueensis* (Figure 01). The high abundance of both host and prophage suggests that these two vOTUs follow a “Piggyback-the-Winner” pattern of viral ecology (Hu et al., 2025). Across all vOTUs, these are the only host matched viruses which directly increase in abundance with corresponding host taxa (Supplemental Material S6).

The only sample site which contains *Gloeobacter* dominated biofilms is the steam pit. The steam wall sample site, located ∼100 m from the pit, is dominated by other oxygenic phototrophic cyanobacteria and is more exposed to light than the shaded steam pit. *Gloeobacter* lacks a thylakoid membrane and is susceptible to light induced stress, resulting in metabolic disruption and cellular damage from oxidative stress (J. H. W. Saw et al., 2013). Our recovery of these vOTUs is the first description of viruses known to infect the *Gloeobacter* genus, which is of astrobiological significance due to its evolutionary role as an ancient cyanobacterium (Prescott et al., 2022; J. H. W. Saw et al., 2013). *Gloeobacter kilaueensis* is a highly conserved species with MAGs sharing 99% ANI (J. H. Saw et al., 2026; J. H. W. Saw et al., 2013). The presence of multiple vOTUs within steam pit biofilms suggests these prophages introduce genetic variation to the *Gloeobacter* host and facilitate genome evolution of the *Gloeobacter kilaueensis* species. CRISPR arrays in *Gloeobacter* MAGs contain a spacer match to a single vOTU of the genus-rank cluster. The match to a single vOTU within the genus could indicate reduced reliance on CRISPR as defense against the prophage or that the host no longer benefits from defense against vOTUs as they diversify in close proximity without geographic partitioning.

In addition to susceptibility to oxidative stress, *Gloeobacter*’s energy inefficiency makes its high relative abundance in biofilms highly unexpected. All membrane-based energy metabolism occurs within a single layer of the cytoplasmic membrane, restricting surface area and metabolic rates. *Gloeobacter*’s reduced photosystems, light sensitivity, and inefficient RuBisCO (slow carboxylation rate) also limit growth of the genus compared to other oxygenic *Cyanobacteria* (Raven & Sánchez-Baracaldo, 2021). Energetically, it is possible that the putative carbon utilization AMG polyhydroxybutyrate depolymerase (*phaZ*) confers host fitness for *Gloeobacter*, contributing to its dominance in steam pit biofilms and optimizing host carbon utilization. Polyhydroxybutyrate depolymerase is an intracellular enzyme which breaks down polyhydroxyalkanoates (PHAs) or polyhydroxybutyrate (PHB) biopolymers. The ability to utilize PHB biopolymers may provide a mobilizable reserve of carbon and energy to *Gloeobacter*, enabling efficient resource management (granules storing nutrients without altering internal osmotic pressure or loss of compounds through leakage) and increasing prophage-mediated *Gloeobacter* survival over other competitors. The vOTU containing the *phaZ* AMG candidate is doubled in abundance compared to the other *Gloeobacter* prophage vOTU, suggesting that there is a comparative fitness benefit to the prophage maintaining the putative AMG.

There is evidence of the Gloeobacter prophage evading host defenses. The presence of what appears to be a self-targeting spacer shows that CRISPR systems of the host may prevent prophage activation or lytic activity (Supplemental Material S7). Alternatively, the prophage may carry an anti-CRISPR protein that inhibits host CRISPR-Cas systems (*Gloeobacter* MAGs contain both Type I and Type III CRISPR-Cas systems). The biofilm sample observed which had both high abundance of *Gloeobacter* (J. H. Saw et al., 2026) and a low abundance of the *Gloeobacter*-linked vOTUs was collected from a sunlight-exposed area, with a lower light intensity than the steam wall sample site. High-light stress in exposed biofilms may compromise *Gloeobacter* cells with metabolic disruption or cellular damage, destabilizing prophage maintenance and reducing their relative abundance in the sample (Figure 01). Oxidative stress could also trigger prophage excision and loss (Middelboe et al., 2025). The substantial decrease of prophage content in biofilms despite the host remaining abundant suggests that light-induced stress in Gloeobacter may limit lysogeny (consistent with “Piggyback-the-Winner” models) influencing viral turnover and potentially altering persistence of lysogenic viral populations.

### Active fumaroles host diverse viral communities

Soil has been described as a medium that contains one of the most diverse and complex viral communities with high richness and evenness suggesting significant levels of viral activity (Roux & Emerson, 2022), In particular, young volcanic soil is known to harbor incredibly diverse microbial and viral communities as a result of the soil’s nutrient rich physicochemical properties (Aguila-Torres et al., 2025; Huang et al., 2024; Mihai et al., 2023). Despite these previous descriptions of high volcanic soil viral richness, we discover that the biofilms of active (thermal and steam emitting) fumarole features contain communities with vOTU richness that is significantly higher than vOTU richness of non-thermal volcanic soil from the Kilauea caldera Big Ell cave (Figure 04). Further, some of the biofilms also contain high vOTU evenness that is comparable to the evenness of volcanic soil. Because biofilms with high viral richness and evenness do not contain a specific microbial phylogenetic profile, we suggest that the environmental context of Hawaiian fumaroles as a geothermal system contributes significantly to the overall viral diversity. There is not a clear distance decay effect or geographic partitioning of diversity indexes, indicating that localized biogeographic context is significant in explaining viral diversity. The structural properties of biofilms and microbial metabolism within fumarole features (increased nutrient availability through basaltic bioleaching, metabolism of emitted gases, and temperature) may all contribute to fumarole viral diversity.

The statistically significant relationship between temperature and viral richness has not been described previously in a geothermal system other than hot springs (Castelán-Sánchez et al., 2020; Sharp et al., 2014). The steady increase of viral diversity in fumaroles which peaks at 50°C and decreases in high geothermal biofilm samples emitting steam (>70°C) provides compelling evidence that viral diversity in some geothermal systems can adhere to a temperature-diversity gradient, and that biodiversity hotspots in geothermal environments are are not limited to “Humboldt’s spa” of hot springs (Sharp et al., 2014). It is known that temperature influences structural biofilm properties, regulating processes such as viral particle attachment and mineral precipitation (Simmons et al., 2020). Indeed, temperature dependent biomineralization has been described within moderately geothermal ranges (Kostešić et al., 2023). These structural alterations could potentially turn biofilms into physical refugia for virion particles, increasing the density of virion particles with immobilization and in turn increasing metagenomically identified viral diversity for species richness while potentially increasing species evenness under specific physicochemical conditions which influence biofilm properties (Decho & Gutierrez, 2017). The physical proximity of viruses within the biofilms exhibiting thermal-induced structural properties could allow these viruses to act as “gene highways” (Thomas et al., 2021) and facilitate microbial adaptation to the dynamic and harsh conditions of a volcanic system (Supplemental Material S8). Even though moderately geothermal samples have high viral richness and evenness and there is a unimodal relationship between temperature and species richness, microbial diversity indexes of samples do not follow the same clear trend (Figure 04).. The decoupling of fumarole virus and microbial diversity could reflect temporal lag and community turnover of viruses, or dormant but inactive viral populations retained at higher abundances than expected within biofilms.

If the viral richness and evenness within biofilms is representative of active viral populations, this may facilitate increased opportunity for viral competition while increasing the fitness conferred by superinfection exclusion (Muscatt et al., 2022; Pinilla-Redondo et al., 2020). This would contribute to explaining diverse and frequent genomic observations of TA systems (encoded in ∼5% of fumarole vOTUs) and other defense related proteins in fumarole viruses (Supplemental Material S8).

### Accessory viral genes may aid adaptation to basaltic surface and resource limitation

Biofilms contain the majority of putative AMGs. Of identified fumarole vOTUs, only 7% of fumarole vOTUs contain AMG candidates which is consistent with observations of AMGs as rare in other geothermal environments such as hydrothermal vents (Langwig et al., 2025). However, more samples with broader geographic distribution are necessary to make conclusions about AMG prevalence in systems with high microbial endemism anticipated (Castelán-Sánchez et al., 2019; Langwig et al., 2025; Salgado et al., 2025; Thomas et al., 2021). The presence of diverse AMG candidates seen within vOTUs suggests that these viruses may function as a genomic reservoir for microbial hosts and facilitate their persistence in adverse and nutrient limited conditions (Supplemental Material S8). Basalt hosted microbial communities in hydrothermal vents are known to engage in bioleaching with enzymatic dissolution of substrate in hydrothermal vents, releasing metals in the process (Bergsten et al., 2021; Thomas et al., 2021). The presence of ten different vOTUs with putative AMG involved in heavy metal detoxification would suggest that viruses play a significant role in facilitating microbial adaptations to high metal concentrations released from the dissolution of basalt (Figure 05). This explains the presence of detoxifying metal efflux and ROS neutralization putative AMGs, which are often seen in hydrothermal vents and heavy metal contaminated soil where similar processes may occur(Cheng et al., 2022; Farias et al., 2015). While most of these transporters are predicted to facilitate heavy metal resistance, some fumarole AMGs candidates likely confer fitness for the host in other ways. MFS transporters may alternatively play a role in conferring antibiotic resistance for the host, while biopolymer transport genes are also observed.

Other AMG candidates increase host fitness in resource limited conditions, primarily with genes that participate in carbon utilization pathways. PHA/PHB metabolism in particular may play an important role in microbial response to adverse conditions. Polyhydroxybutyrate depolymerase is an AMG candidate in a *Gloeobacter* prophage vOTU which is highly abundant in steam pit samples (Supplemental Material S8). Two other vOTUs contain carbon utilization putative AMGs which could be involved in either beta oxidation of fatty acids or PHA biosynthetic pathways (conversion of short-chain carboxylic fatty acids). In resource limited conditions with nutrient depletion, PHA and PHB utilization could provide microbes the ability to sequester and mobilize carbon and energy sources, aiding survival in dynamic and volcanism dependent physicochemical conditions of fumaroles. Chitin degrading genes (chitinase and chitosanase) are contained by three different fumarole vOTUs and are AMG candidates. Chitinase has been described as an AMG in prophage encoded chitinase within hydrothermal vents, where the AMG is involved in substrate dependent lysis-lysogeny switches of viruses. It is known that carbohydrate degradation capacity is a driver of microbial community structure in hadal zones, offering a possible explanation for how these putative AMGs could confer host fitness in nutrient limited and dynamic environments such as fumarole biofilms (Middelboe et al., 2025). Putative AMGs related to nutrient limitation suggest viral AMGs confer host fitness with AMGs related to resource acquisition and scavenging (Monier et al., 2012; Tian et al., 2024; Zheng et al., 2021).

Sporulation related AMG candidates may facilitate survival in adverse conditions via dormancy, which has primarily been observed as an AMG in low oxygen zones, heavy metal contaminated soil, and sub-freezing environments (Brown et al., 2022; Jurgensen et al., 2022; Sun et al., 2023). These putative AMGs may help their host by facilitating microbial sporulation when conditions are resource limited until nutrients are available for host reactivation. The selection for microbial survivability under resource limitation may incentivize acquisition of viruses with these AMG candidates. However, identification of viral metabolic genes is constrained by sequence length (contigs under 10 kb). Biochemical characterization or transcriptomic changes provide evidence for confident AMG identifications, and should be performed for any structurally novel predicted AMGs. This is particularly true for CAZyme genes which are poorly conserved on a structural level (Martin et al., 2025).

## Methods

### Sample collection and sequencing

Detailed methods on sample collection and sequencing have been described in two previous studies (Prescott et al., 2022; J. H. Saw et al., 2026). Briefly, a total of 46 biofilm and soil samples were collected in an expedition in 2019 from steam vent features found within Kilauea Caldera inside Hawaii Volcanoes National Park and within East Rift Zone along Hawaii Route 130. DNA from samples were extracted using ZymoBIOMICs mini prep kit, quantified with Qubit HS dsDNA assay, then sent for sequencing at two sequencing centers: ASGPB at University of Hawaii at Manoa for 16S amplicon sequencing and CQLS at Oregon State University for whole-genome shotgun (WGS) sequencing using Illumina MiSeq and HiSeq, respectively. Some of the amplicon data has been analyzed in a previous study that excluded the cave soil samples from Big Ell (Prescott et al., 2022). BBTools script bbduk.sh (Bushnell, 2014) was used for quality control prior to assembly. Adapters were trimmed from the right ends of the reads, with reads shorter than 50 bp discarded (--minlen=50). Adapters were trimmed based on paired overlap (tbo=t) with low quality bases trimmed on both ends of reads (--qtrim=rl) with low quality bases discarded (--trimq=20).

#### Identifying viral sequences

The ViralAssembly module of metaviralSPAdes v3.15.5 (Antipov et al., 2020) was used for viral assembly and mismatch correction (k-mer lengths of 21, 33, 55, and 77). Assembled contigs were input into ViralVerify v1.1.0 (Antipov et al., 2020) (Naive Bayesian classifier on hmmsearch gene predictions), geNomad v1.11.2 (Camargo et al., 2024) (hybrid approach of alignment-free models and gene models with neural networks for classification), and VirSorter2 v2.2.4 (Guo et al., 2021) (multi-classifier with HMM search for viral hallmark genes) for initial identification. These tools employ a diversity of methods for viral identification (Antipov et al., 2020; Bolduc et al., 2025; Camargo et al., 2024; Guo et al., 2021). Viruses were then identified by identifying both a head and packaging gene in each candidate viral contig. Default parameters were used for ViralVerify with the ‘-hmm’ flag. geNomad was run with the ‘--end-to-end’ flag. As per community standards for VirSorter2, the default score of 0.5 (low-cutoff) was used for maximum sensitivity and filtering. CheckV v1.0.3 (Nayfach et al., 2021) was then used on extracted contigs (end_to_end option) to remove false positives and trim host regions of proviruses prior to running VirSorter2 again on trimmed sequences (Guo et al., 2021; Tian et al., 2024). Lastly for the VirSorter2 pipeline, a secondary screening approach was employed on domain architecture DRAM-v annotations searched for capsid associated proteins (Phage-coat PFAM clans CL0373 or CL0371). PFAM clans account for evolutionary relationships which cannot be encompassed by a single HMM profile of DRAM-v (Finn et al., 2016; Kashif-Khan et al., 2024).

Open reading frames (ORFs) were called in metagenomic (‘-meta’) mode with Prodigal v2.6.3 (Hyatt et al., 2010). Gene calls were annotated with HHblits (Steinegger et al., 2019) (uniprot_sprot_vir70 database) with flag ‘-n 3’ iterations for remote homology prediction. Annotations were filtered for E value (<1E-1) and probability (>50%). The first 10 hits for each Coding DNA Sequence (CDS) were screened for head and packaging associated genes using an automated python script which iterated over annotations by contig. A list of 33 gene ontology (GO) terms identified as head or packaging genes was curated from the QuickGO display (Binns et al., 2009). UniProtKB annotations which correspond to GO terms for virion assembly and maturation were extracted and curated to exclude baseplate and tail gene products (The UniProt Consortium, 2022). The resulting list of 332 UniProtKB codes were used for the Python search script which flagged contigs containing both head and packaging associated gene products which were then manually verified. All manual checks for gene functional prediction were performed with a combination of BLAST(Camacho et al., 2009), UniProt BLAST, and HHPred (Swissprot DB) (Söding et al., 2005).

### Viral clustering

Viral sequences were also grouped into roughly genus level viral clusters (VCs) using vConTACT3 v.3.0 with database (v220) domain restricted to “prokaryotes” (Bolduc et al., 2025). Graph output (.cyjs) was imported into Cytoscape (v3.10.2) and the NetworkAnalyzer plugin was used to evaluate the quality of clusters (Su et al., 2014). ‘Clustering Coefficient’ and ‘Topological Coefficient’ metrics were filtered to be equal to or greater than 0.85 (Tian et al., 2024). Subclustering (virus cluster of three or more nodes without links to the rest of the network) is observed for 20 VCs. Packaging and head related genes of viruses that were unable to be taxonomically assigned into a realm by vConTACT3 v.3.0 were manually verified using BLAST, UniProt BLAST, and remote homology prediction tool HHPred. Genes for major capsid protein and TerL subunit were annotated (HHblits) and clustered (BLAST 2.14.1+) to assess cluster success with viral sequences of variable length (variation > 1 kb).

### Generating vOTUs

Viruses were clustered into species-rank vOTUs reflecting viral populations using Minimum Information about an Uncultivated Virus Genome (MIUViG) standards with recommended thresholds (95% ANI and 85% AF) for clustering parameters (Roux et al., 2019). Contigs were first compared with nucleotide BLAST (v2.11.0+) and pairwise comparisons were generated by blasting all sequences together (“-outfmt ‘6 std qlen slen’ -max_target_seqs 10000”). Finally, clustering of viral sequences was carried out using scripts anicalc.py and aniclust.py (“--min_ani 95 --min_tcov 85 --min_qcov 0”) which are distributed as part of the CheckV package (Nayfach et al., 2021).

### Calculating vOTU diversity and abundance

Individual viral contigs were mapped using Bowtie2 (Langmead & Salzberg, 2012) with the bowtie2-build function to create .sam files. The .sam files were converted to .bam files then sorted, indexed, and counted (samtools idxstats) with SAMtools (Li et al., 2009). Relative abundance was calculated with reads per kilobase per million mapped reads (RPKM) for each viral contig and then summed for all contigs for each vOTU (Yan et al., 2025). vOTU richness and evenness were calculated with the vegan package v2.7-2 (Oksanen, 2025) in R. Richness was assessed for each sample by finding the Chao1 index using the sample vOTU relative abundance table, which was calculated with the specnumber() function. Shannon diversity was calculated for each sample with the diversity() function from the sample relative vOTU abundance table. Pielou’s evenness was calculated by dividing the Shannon index by the ln() of Chao1. The relationship between the temperature of samples and vOTU diversity indexes was assessed for significance with a non-parametric least squares regression, and then assessed for overfitting with a Bayesian information criterion test with sample correction (BICc) due to sample size (n=20). Linear, quadratic, and cubic models were attempted, and the lowest BIC value indicated the best fit model. Normal distribution and constant variance of residuals was supported with density and QQ-plots for both relationships (Supplemental Figure 07). Normality for models of temperature and related diversity indexes was confirmed with the Shapiro-Wilk test.

### Phylogenetic classification of viruses

Phylogenetic estimation was guided by vConTACT3 v.3.0 (Bolduc et al., 2017, 2025). Phylogenies were estimated using maximum likelihood independently for classes Caudoviricetes, Malgrandaviricetes, Laserviricetes, and Tectiliviricetes. Caudoviricetes references were chosen by selecting first neighbors (‘first neighbors of select nodes’ Cytoscape v3.10.2) of vConTACT3 v.3.0 assigned fumarole Caudoviricetes (Shannon et al., 2003). For Malgrandaviricetes, Laserviricetes, and Tectiliviricetes phylogenies, all known references were selected from ICTV MSL40.v1 and ORFs were called with Prodigal v2.6.3 (Hyatt et al., 2010). All references were annotated as described above with HHblits to identify marker genes. The marker genes were TerL, major capsid, and packaging ATPase marker genes for the Caudoviricetes, Malgrandaviricetes, Laserviricetes,Tecitiviricetes trees respectively with 1000 iterations of tree topology, branch lengths, and substitution model parameters (Grybchuk et al., 2024; Katoh et al., 2002). Trimming of the multiple sequence alignment was performed with trimAl v1.5.rev0 with a gap threshold of 30% (Capella-Gutiérrez et al., 2009). IQTree v2.4.0 was used for phylogenetic reconstruction, with the evolutionary model Q.pfam+F+I+G4 selected as the best model for *Caudoviricetes* (Nguyen et al., 2014). The model Q.pfam+G4 was used for Malgrandaviricetes and *Laserviricetes* trees, and the Q.pfam+I+G4 model was used for *Tectiliviricetes*. 1000 bootstrap estimates were used for branch support estimation. All phylogenies were midpoint rooted and visualized with iTOL v7 (Letunic & Bork, 2006).

### Host matching with CRISPR spacers

Putative host prediction was performed with CRISPR spacer matching. CRISPR-Cas systems were identified from the binned nucleic acid sequences with <5% contamination using the standalone tool of CRISPRCasFinder as a Singularity image (Couvin et al., 2018). High confidence arrays were identified by using ‘-levelMin 3’ parameter. Arrays adjacent to Cas systems within 10,000 base pairs (bp) were identified with the ‘-ccc’ parameter. Spacers were extracted from high confidence arrays using minCED and clustered with a cutoff of 100% identity using cd-hit-est (directionality assessed with parameter ‘-r 1’) to dereplicate spacer sequences (Bland et al., 2007; Li & Godzik, 2006). Targets with host matches were sorted into three categories of mobile elements during the viral identification workflow: plasmid, viral, or unknown mobile genetic element (MGEs). MGEs that did not have chromosomal, plasmid, or viral indicators (structural genes) were characterized as other MGEs. Chromosomal fragments were confidently excluded by scanning matches and eliminating targets with either entire arrays or multiple shared spacers and neighboring sequences which were marked as ‘suspicious’ matches (Supplemental Material S7). Viruses identified as viral and plasmid hits or uncertain viral and plasmid sequences (ViralVerify, VirSorter2, and geNomad) were aligned with BLAST (2.14.1+). Matches were filtered for ratio *nident* (identical bp matches) to *qlen* (spacer length) > 0.89 to be considered operationally valid.

Matches that could be plasmid-encoded arrays or chromosomal fragments were eliminated as follows (Pinilla-Redondo et al., 2021): CRISPRDetect was used to identify arrays in chromosomal arrays and target elements which were then aligned with BLAST (2.14.1+). Identical matches between chromosomal and plasmid arrays were discarded immediately. If there were multiple shared spacers and neighboring sequences (spacers and repeat element) the hit was designated as suspicious (Supplemental Material S9). Target sequences were searched for ORFs using Prodigal (v2.6.3) and .faa (amino acid sequence) and .sco (ORF coordinates) output files were used in conjunction with BLAST (2.14.1+) match coordinates to identify which ORF was targeted by spacers. Targeted ORFs ( coding sequences (CDS) or non-coding ORFs) were annotated using BLAST, UniProtBLAST, and HHPred.

#### Viral genome annotation (metabolic and accessory genes)

Three categories of genes were systematically recovered from vOTUs: auxiliary metabolic genes (AMGs) that may have a role in host metabolism during the infection cycle, viral resistance genes, and viral defense system related genes. Viral resistance and defense genes are non-essential accessory genes (morons) which provide an adaptive advantage to either the phage or host (Kang et al., 2024). Annotation for functional gene product prediction was performed with DRAM-v v0.1.2 (Shaffer et al., 2020) on all viral sequences. Final vOTUs were processed with VirSorter2 v1.0.1 using the -prep-for-dramv flag then annotated with DRAM-v to identify viral metabolic genes. Gene curation was performed by filtering and manually scanning both the DRAM-v AMG summary (275 gene predictions) and total annotation output (20,666 genes) for AMGs and accessory genes. Gene product annotations were discarded for genes that did not have an auxiliary score <4, or had DRAM-v auxiliary flags of (V) viral structural component, (A) attachment, (P), viral specific MEROPS peptidase, or (F) ‘near end of contig’ (ter Horst et al., 2021; Tian et al., 2024). To eliminate the candidate genes within host-like gene operons (possible miscalled proviral ends) all candidate genes were scanned for flanking viral genes using DRAM-v flag (V) for structural virion components, annotations such as portal, tail, baseplate, capsid, and structural or replicative genes with VOGDB flags (Xs;Xr). This left 212 putative AMG candidates (Supplemental Material S8).

To be conservative in AMG prediction, secondary manual curation was performed on putative AMG candidates to eliminate genes that could be involved in viral life cycle processes (ter Horst et al., 2021; Trubl et al., 2018). This led to removal of putative AMGs involved in nucleotide or amino acid metabolism (DRAM-v and KEGG-MAPPER pathways and modules). Putative AMGs were also discarded if their annotation was overly general (genes annotated as glycosyltransferases, peptidases, glycosylhydrolases, adenylyltransferases, methyltransferases, and ribosomal proteins). Attempts were made to obtain a more specific gene prediction with BLASTp before discarding (Martin et al., 2025; ter Horst et al., 2021). To predict accessory genes, a secondary manual curation was performed by searching non-metabolic annotations that did not have the (M) auxiliary flag in the DRAM-v master annotation spreadsheet output. Participation in life cycle pathways was assessed using methods described above, and resulting accessory gene candidates were manually inspected.

Prophage annotation and boundaries were performed by aligning viral contigs to MAGs with BLAST+ alignment (2.14.1+). Prophage boundaries were identified in viral contigs PHASTEST and manually checked for flanking viral genes on viral contigs using annotation methods described in the workflow for viral identification (Wishart et al., 2023). ORFs were called with Prodigal (v2.6.3) and annotated with Pharokka (v1.7.5) identifying phage orthologous groups (PHROGs) with local alignment (Bouras et al., 2022). Gene diagrams (genomic context determined with MMSeq2 and pyHMMer) were plotted using LoVis4u (Egorov & Atkinson, 2025; Steinegger & Söding, 2017). Proteomic similarity matrix for *Microviridae* viruses was built with MMSeq2 alignments through LoVis4u.

## Acknowledgements

We would like to thank George Washington University’s high-performance computing facility, Pegasus, for providing data storage, support, and computing power for metagenomic analyses. We would also like to thank Pietro Tardelli Canedo, Pooja Anilkumar, Arash Raeisbahrami, Zachary J. Watts, and Max Seledes for discussion, as well as the International Viromics

Workshop at Ohio State University which provided authors Pia Sen and Lausanne Lee Oliver foundational knowledge for viral metagenomic analysis. We also thank Keith A. Crandall, Sarah Johnson, and Gary Trubl for manuscript review and comments. We gratefully acknowledge the computing resources provided on the High-Performance Computing cluster operated by Research Technology Services at The George Washington University. K.S.M. and Y.I.W. were supported by the Intramural Research Program of the National Institutes of Health (NIH). The contributions of the NIH authors are considered Works of the United States Government. The findings and conclusions presented in this paper are those of the author and do not necessarily reflect the views of the NIH or the U.S. Department of Health and Human Services.

## Data availability

All raw data used in this study have been deposited in the European Nucleotide Archive (ENA) under project accession number PRJEB52128. Phylogenetic IQ-Trees, viral candidate contigs, and final viral contigs are available on Zenodo at DOI: https://doi.org/10.5281/zenodo.19378070

## Code availability

Scripts to run analyses and viral identification pipeline custom scripts are available at https://github.com/pia-sen/Hawaii_fumarole_viruses

## Author contributions

J.H.S. designed and led the sampling and sequencing. P.S. and J.H.S. determined the scope of analysis for this project. P.S, L.O, C.P. and M.S. performed data analyses, K.S.M. and Y.I.W. supervised and guided development of the viral identification pipeline, P.S. L.O. and J.H.S wrote the manuscript, and all authors reviewed and approved the final version of this paper.

Sequencing of the samples was made possible by the startup funds provided to J.H.S by The George Washington University.

## Supplemental Figures

**Supplemental Figure 01.**
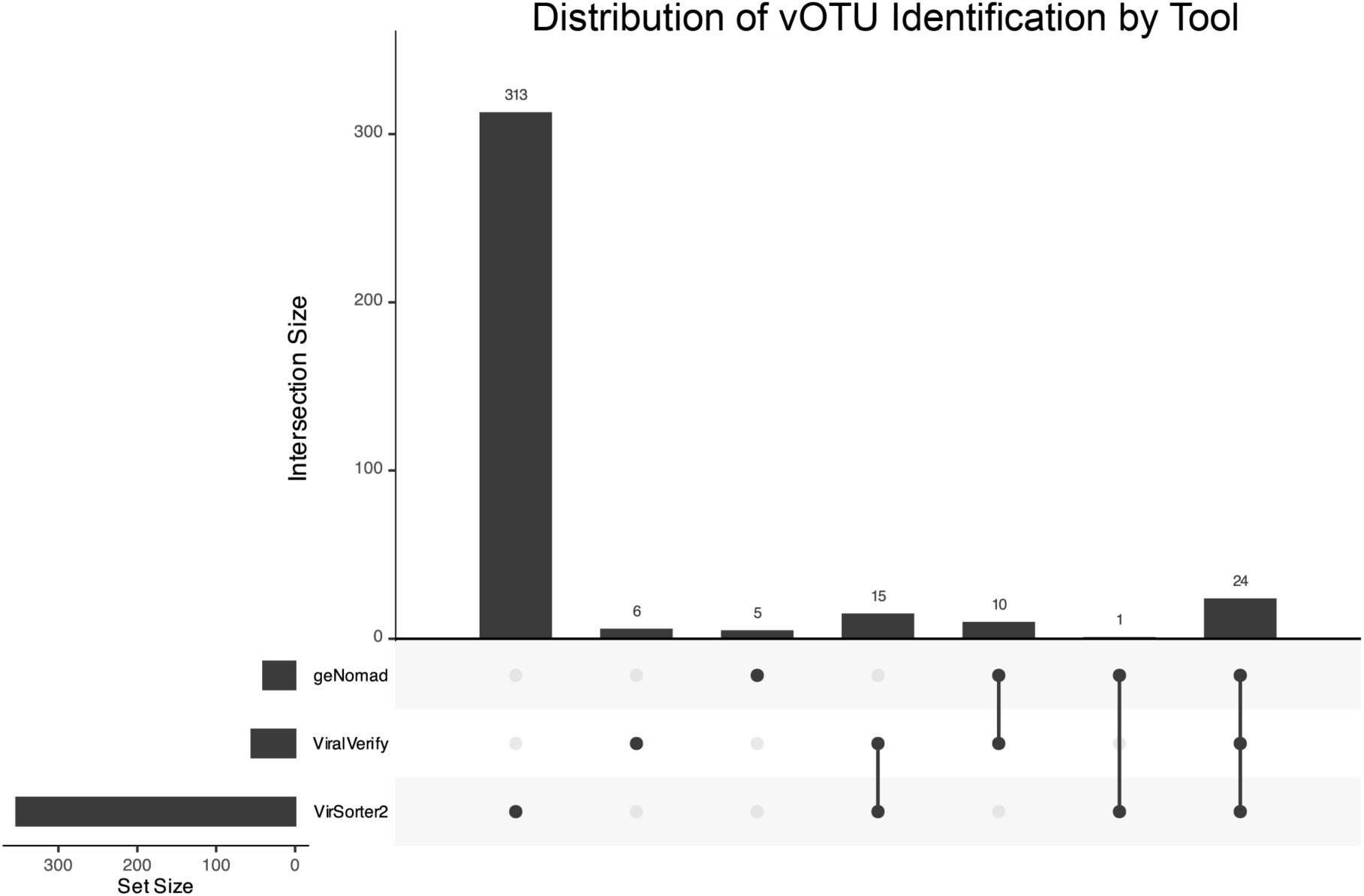
Upset plot of vOTUs identified by each tool. Bars (with counts above) show the number of vOTUs shared across tool combinations indicated by connected, filled points below. The left panel shows total vOTUs identified per tool.

**Supplemental Figure 02.**
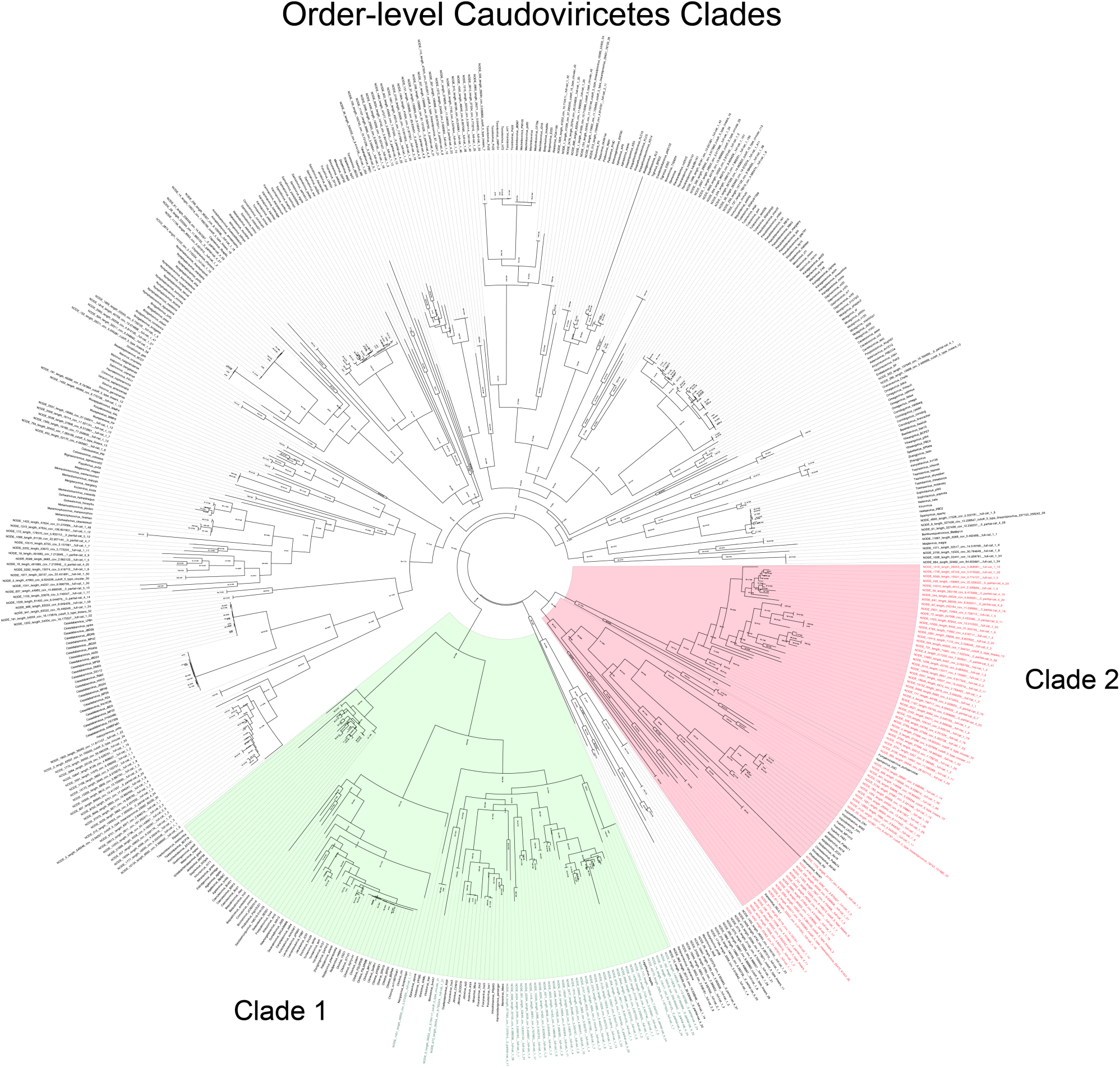
Maximum likelihood tree of Caudoviricetes (TerL) amino acid sequences showing two previously undescribed order-level clades. Clade 1 (Kilaueavirales) is highlighted in green, with green taxon labels indicating fumarole-derived viruses and black taxon labels representing RefSeq viruses (first neighbors). Clade 2 (Pahoavirales) is highlighted in red, with red taxon labels indicating fumarole-derived viruses and black taxon labels representing RefSeq viruses. Node support values are shown and available in the IQ-TREE files on Zenodo.

**Supplemental Figure 03.**
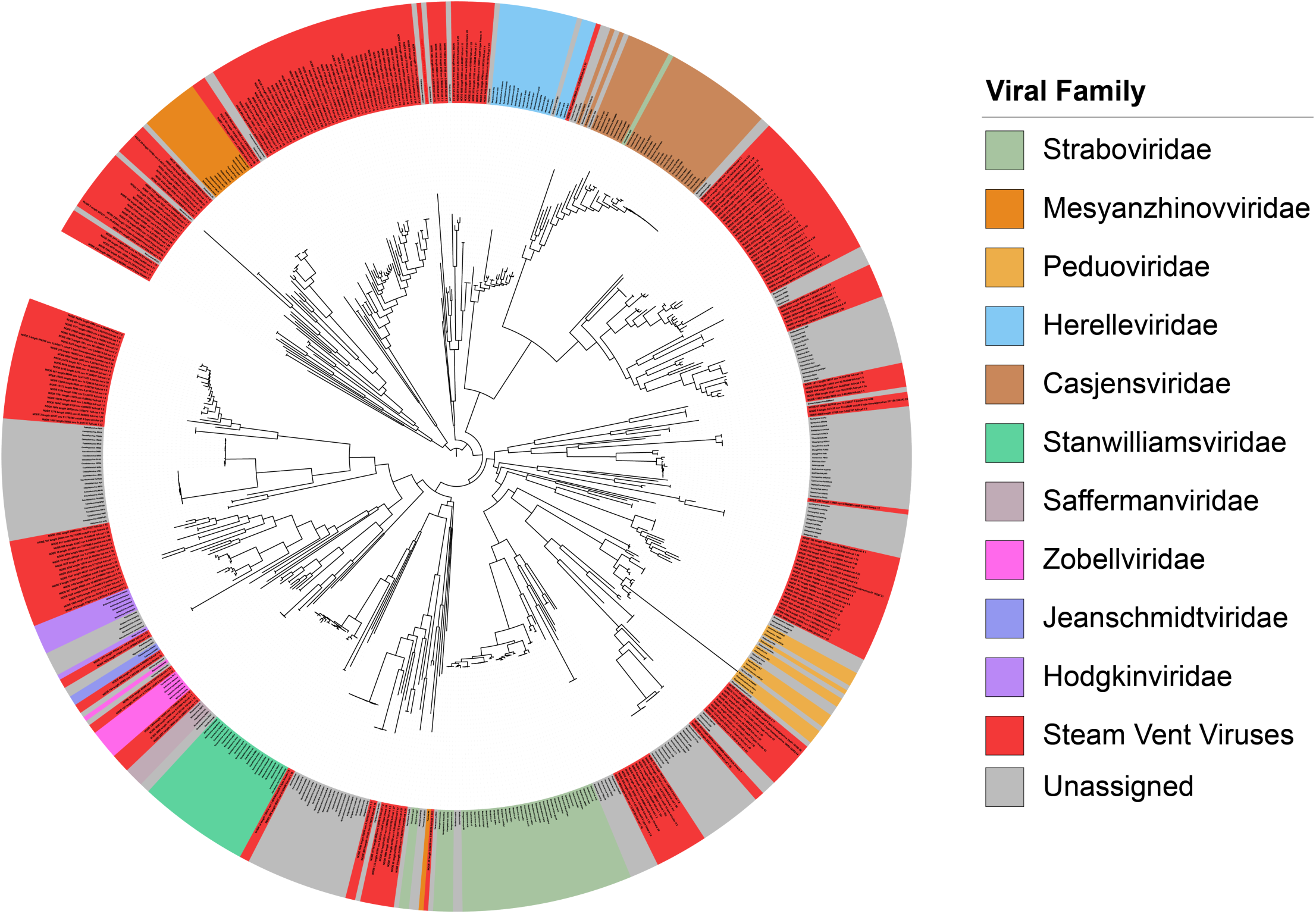
Maximum likelihood tree of Caudoviricetes (TerL) amino acid sequences showing relationships between fumarole viruses and previously described viral families. Node label colors (outer ring) indicate viral family classification (many lacking order-level assignment); grey denotes unassigned viruses, and red denotes steam vent virus TerL. Clade 1 (Kilaueavirales) groups with RefSeq genomes from two families lacking prior order assignment—Casjensviridae and Herelleviridae—and includes 30 fumarole vOTUs and 17 unassigned RefSeq viruses. Clade 2 (Pahoavirales) includes 56 fumarole vOTUs and places Mesyanzhinovviridae as more recently diverged relative to some fumarole viruses. Additional fumarole viruses show close phylogenetic similarity to incertae sedis families, including Peduoviridae, Zobellviridae, and Saffermanviridae.

**Supplemental Figure 04.**
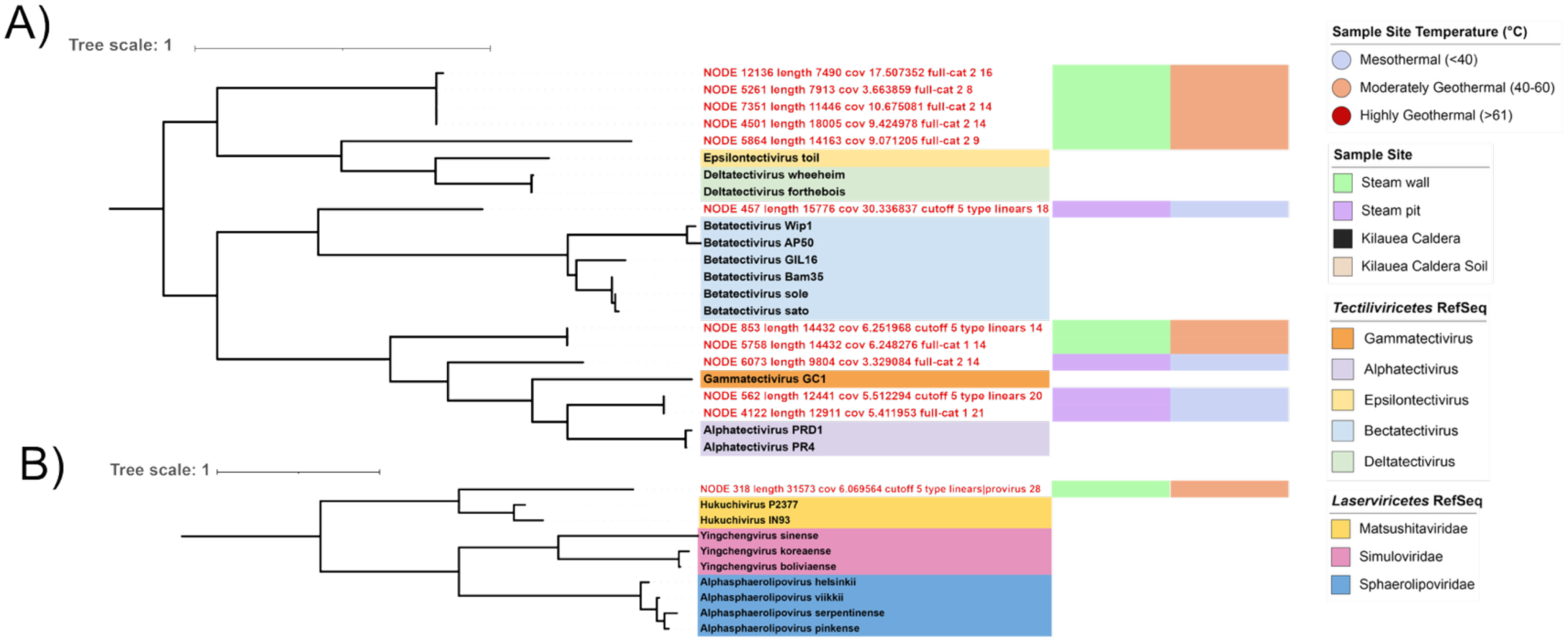
**A)** Maximum likelihood class-level tree of Tectiviricetes packaging ATPase amino acid sequences (annotated with HHblits). Red labels indicate fumarole-derived viruses, while Tectiviricetes RefSeq viruses are highlighted with genus-specific colors as described in the figure legend. Alphatectivirus is the only genus within Tectiviricetes containing plasmid-dependent phages, and all five genera are included as references. **B)** Maximum likelihood class-level tree of Laserviricetes packaging ATPase amino acid sequences (annotated with HHblits). RefSeq viruses are colored by genus as described in the figure legend, and the red label denotes a fumarole-derived virus closely related to Hukuchivirus, a genus of thermophilic Thermus phages.

**Supplemental Figure 05.**
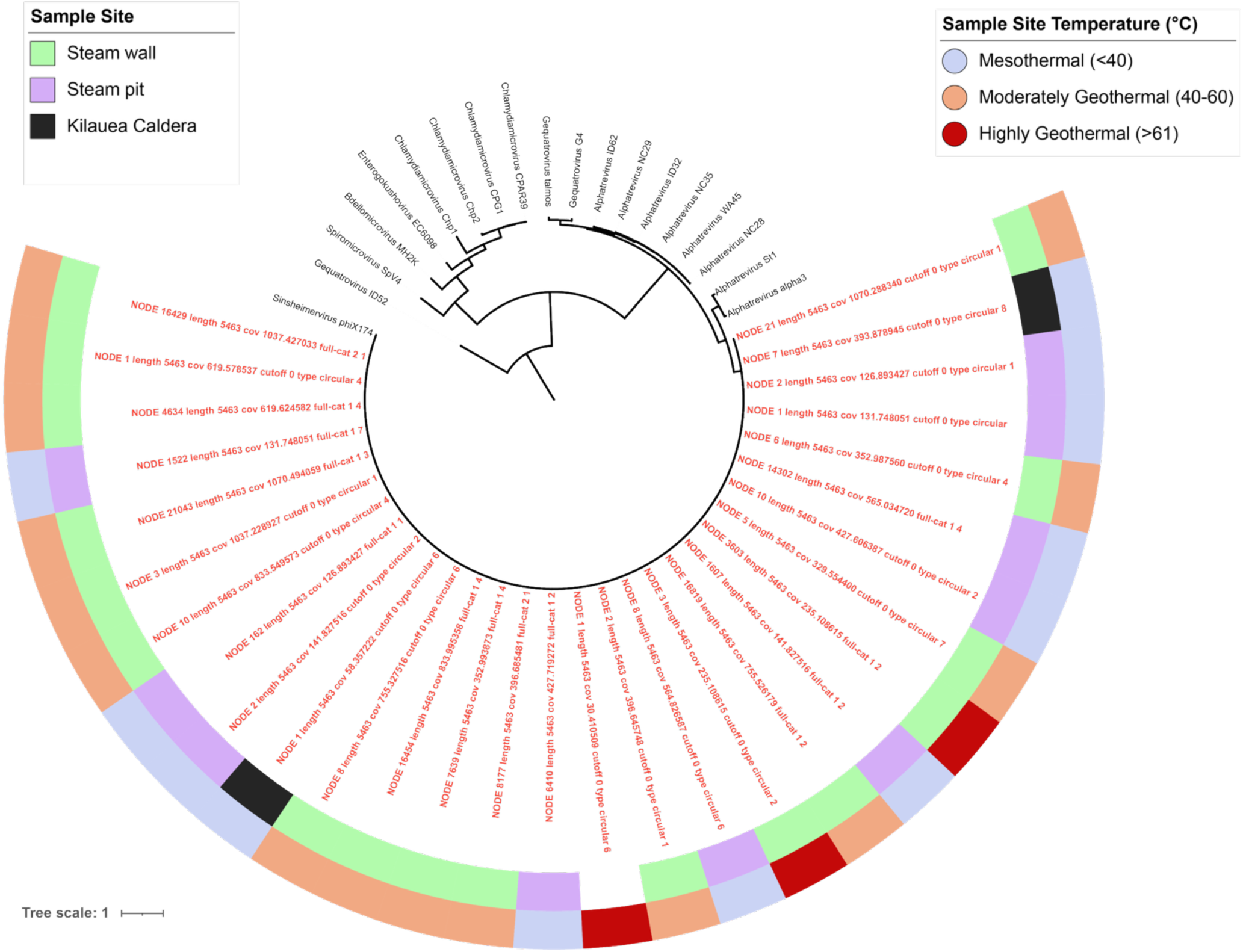
Maximum likelihood class-level tree of Malgrandaviricetes major capsid protein (MCP) amino acid sequences. RefSeq viruses from a shared cluster in the gene-sharing network (whole genomes) were used as references for phylogenetic estimation. Red labels indicate fumarole viruses. The outer ring denotes the temperature range of the source sample, and the middle ring indicates the corresponding sampling site. Fumarole viruses form a poorly resolved clade with phiX174.

**Supplemental Figure 06.**
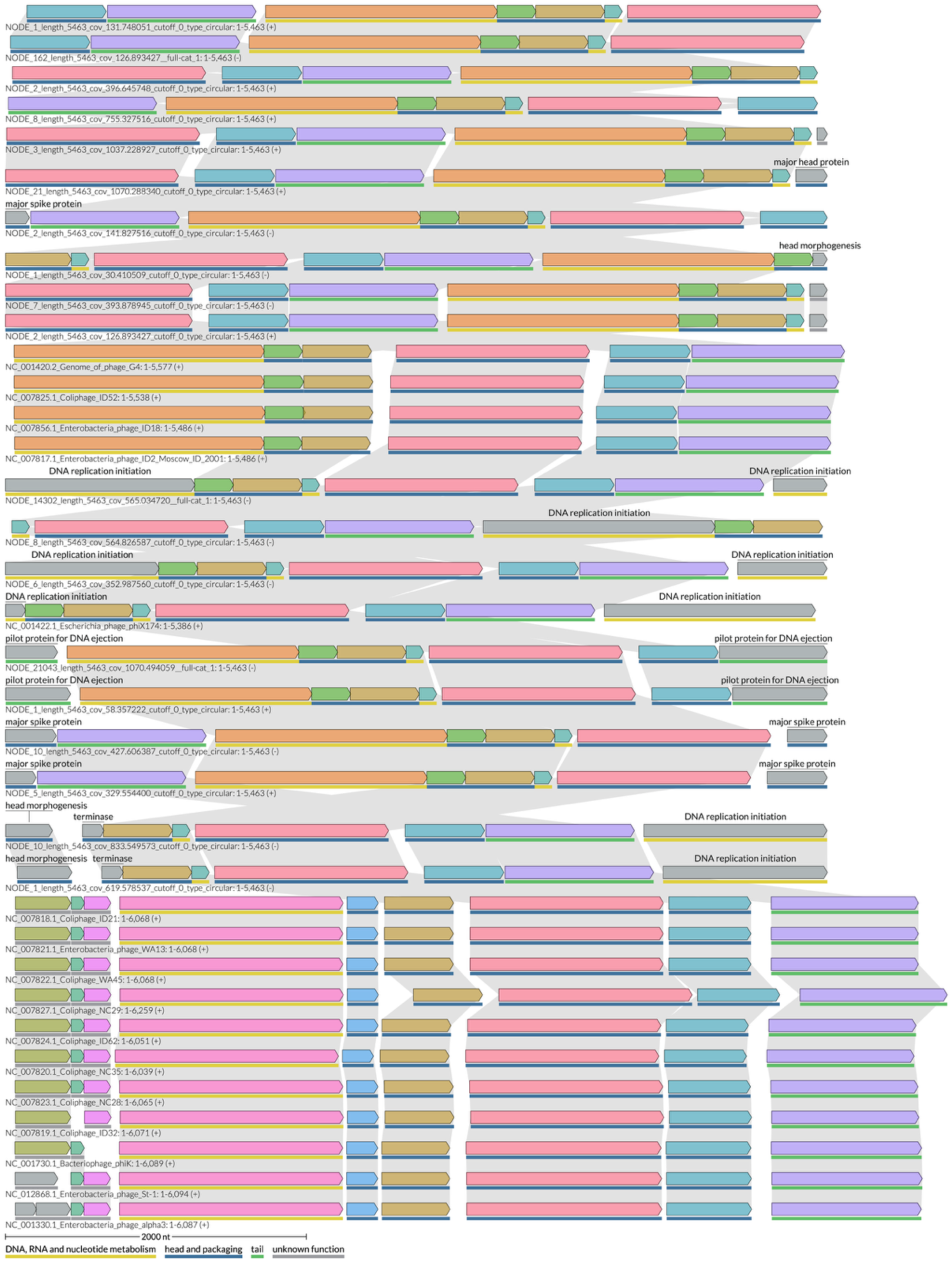
Whole-genome comparative map of Microviridae viruses within the Bullavirinae (phiX-like) subfamily. Genome names are shown to the left below each gene map, with corresponding NCBI accessions when available. Genes are depicted as blocks, with shared colors indicating conserved proteins. Conserved and variable genes are identified using a pairwise proteome composition distance matrix (LoVis4u), and syntenic rearrangements are indicated by linkages between genomes. The bar beneath each genome shows functional annotations from LoVis4u based on local alignments (MMSeq2). Compared to RefSeq Bullavirinae (phiX-like viruses), fumarole Microviridae display greater syntenic variation and distinct genomic differences both from RefSeq genomes and among themselves.

**Supplemental Figure 07.**
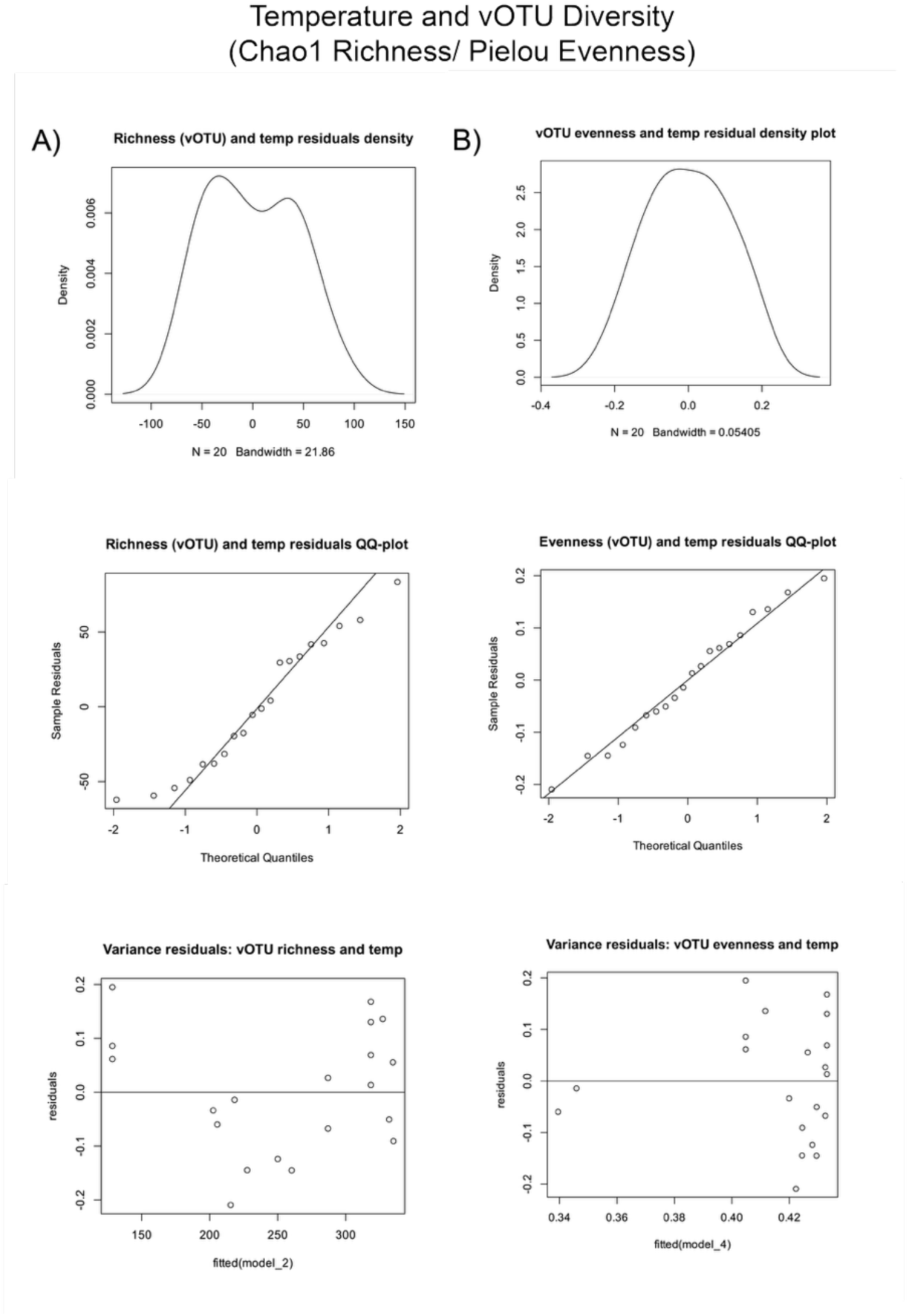
A) Diagnostic plots for the non-linear regression of temperature and vOTU richness (Chao1), including density, QQ, and variance plots of residuals. Residuals show a bimodal distribution in the density and QQ-plots, with constant variance. This supports the use of ordinary least squares (OLS) regression to assess the relationship between temperature and vOTU richness. B) Diagnostic plots for the non-linear regression of temperature and vOTU evenness (Pielou), including density, QQ, and variance plots of residuals. Residuals appear normally distributed with heteroscedasticity, indicating non-constant variance; thus, least squares regression is not sufficient to model the relationship between temperature and vOTU evenness.

**Supplemental Figure 08.**
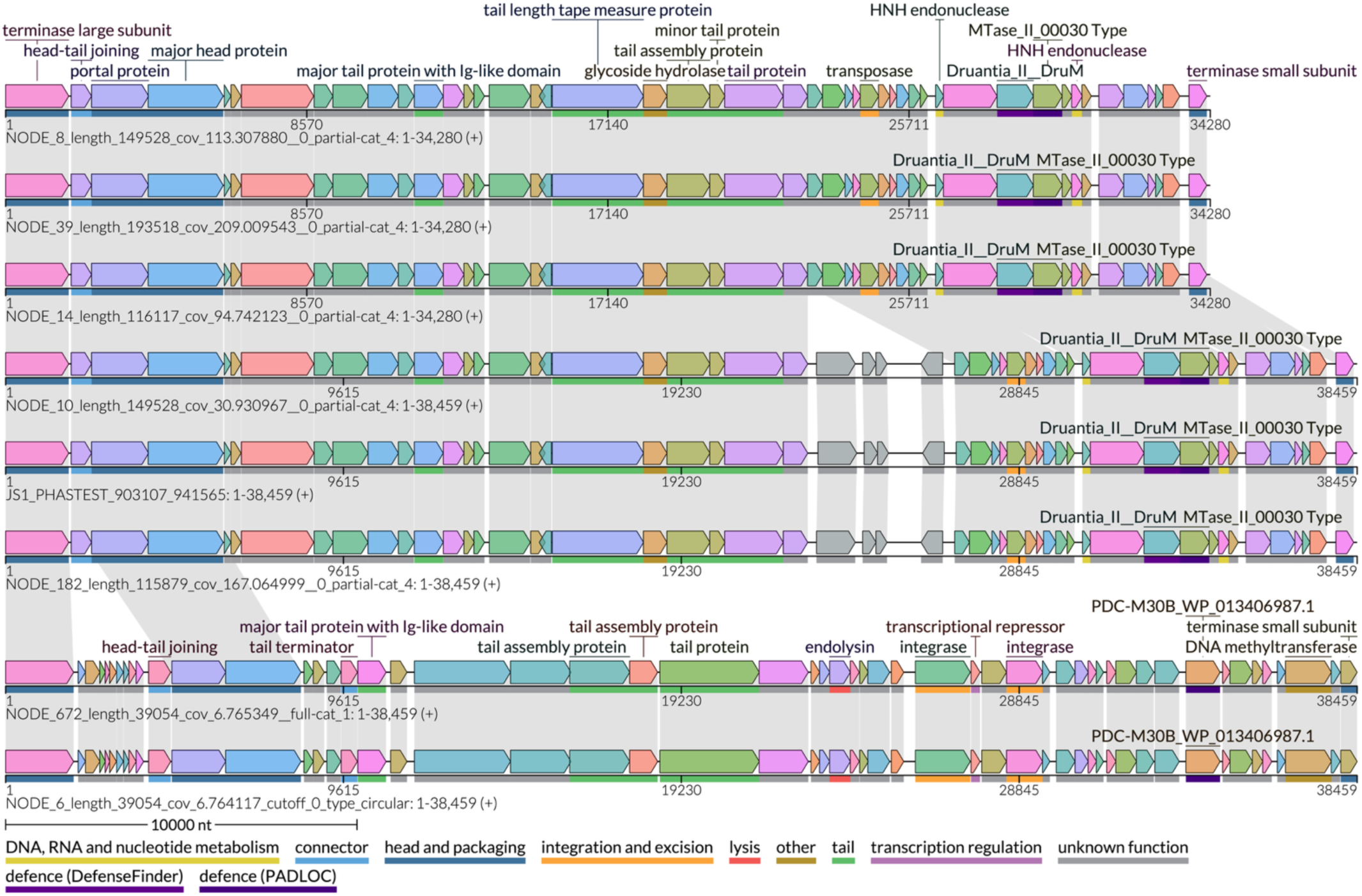
Comparative genomic map of three fumarole vOTUs within a vConTACT3 genus-level cluster. These vOTUs were detected in high abundance only at the steam pit site; two vOTUs show CRISPR-spacer matches to Gloeobacter kilaueensis, representing the first report of a virus infecting this genus. Genome names are shown to the left below each map, with NCBI accessions where available. Genes are depicted as blocks with shared colors indicating conserved proteins. Conserved and variable genes were identified using a pairwise proteome composition distance matrix (LoVis4u), with syntenic rearrangements indicated by linkages. The bar beneath each genome shows functional annotations from LoVis4u based on local alignments (MMSeq2).

